# Proximity labelling of D1-like dopamine receptors reveals distinct cellular environments and uncovers trafficking proteins that regulate DA mediated behaviors in *Drosophila*

**DOI:** 10.64898/2026.05.28.728438

**Authors:** Dana C. Guhle, Bhavana Kanagala, Rebecca Dust, Ross Evashkevich, Ronald L. Davis, Jacob A. Berry

## Abstract

The neurotransmitter dopamine (DA) is central to synaptic regulation that support diverse behavioral functions, including both learning and forgetting. This multi-functional role of DA is due to receptor specific signaling in specific subcellular environments that remain uncharacterized. Here we utilized proximity labelling proteomics in human cells to characterize the proximal environments of two *Drosophila* D1-like DA receptors (Dop1R1 and Dop1R2) in basal and DA activation environments. While DA drives both receptors to recruit Beta-Arrestin 2, Dop1R1 alone showed ligand driven recruitment of G-protein Receptor Kinase 2/3, proximity to clathrin mediated endocytosis, and WASH complex mediated endosomal trafficking. Additionally, we show evidence that Dop1R1 and Dop1R2 reside in distinct domains at the cell surface. *In vivo* disruption of *Drosophila* orthologs of Dop1R proximal proteins revealed three trafficking proteins, Sec24AB, Krz, and CG13887, that regulate R1-mediated learning, starvation induced attraction to odors, and DA-mediated cAMP responses in memory circuits. In addition to revealing DA receptor trafficking proteins that support learning, our comparative characterization of the cellular environments D1-like receptors offers insights into how DA differentially regulates diverse behavioral and synaptic functions.

**For TOC only:** 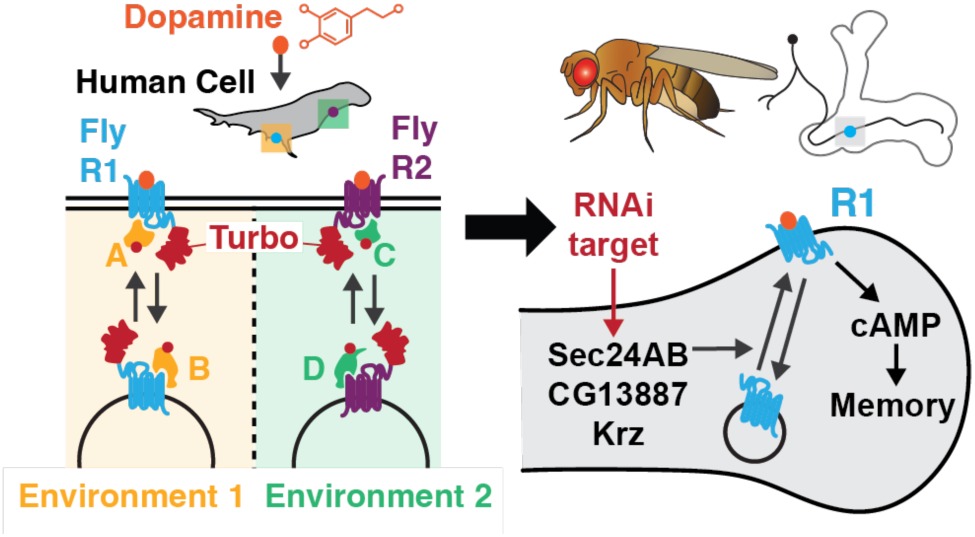

## Introduction

The ability for animals to adapt behaviorally to their environment is a critical brain function that evolved for survival in an ever-changing environment. Synaptic plasticity, the ability to strengthen or weaken connections between neurons, is essential for learning and adaptive behaviors. Across the animal kingdom, the neurotransmitter dopamine (DA) plays a key role in synaptic regulation that underlies adaptive behaviors, including learning^1–4^. Interestingly, DA directs distinct cellular signaling pathways and alters synapses through distinct dopamine receptors in mammals^5,6^ and insects^7–11^. This multi-functionality of DA occurs from receptor-specific interactions occurring in local cellular environments called nanodomains^12–14^. Recently, proximity labelling has emerged as a unique way to covalently label the local environments (∼ 10 nm) around a protein of interest^15–17^. Proximity labelling has been used with a few GPCRs with ligand-binding experiments^18^, or with a single dopamine receptor (D2-like)^19^. However, a comprehensive understanding of the subcellular environments of two similar but distinct dopamine receptors is lacking and critical to understanding the multiple functions of DA.

In *Drosophila*, DA is central for memory encoding and storage in the mushroom body (MB) memory center of the brain^1,2,20–24^. Two D1-like DA receptors Dop1R1 (R1) and Dop1R2 (R2) are highly expressed in the principal neurons of the MB (referred to as MBN)^11,25^, and play a role in distinct aspects of memory, including R1 signaling for the formation of short-term memory^2,26^ and R2 signaling for forgetting^27,28^, long-term memory^29,30^, and memory flexibility^8^. Beyond associative memory, R1 and R2 also play distinct roles in other forms of behavior, including R1-mediated innate odor attraction in hungry flies^31^, and distinct roles in sleep^32^. When R1 and R2 are expressed in human HEK293T cells, they have distinct G-protein coupling and downstream secondary messenger signaling^7^. Specifically, DA activation of R1 drives strong Gas coupling and cAMP generation, whereas activation of R2 weakly couples to Gas and cAMP, but uniquely and robustly couples to Gaq to generate Ca^2+^ release from the endoplasmic reticulum (ER). Remarkably, this cellular signaling is conserved within native contexts in the MBN synapses *in vivo*, and both cAMP (via R1) and Ca^2+^ released from the ER (via R2) are critical and differential regulators of MBN synaptic strength, memory encoding, and forgetting^7,8,27^. However, characterization of the distinct interactomes and local environments before and during DA activation of these two receptors is lacking.

In this study, we utilized TurboID-based proximity labelling and quantitative proteomics in cell lines to comprehensively characterize the proximal environments and potential interactors of two *Drosophila* D1-like DA receptors (R1, R2), in both basal and DA activation environments. We offer evidence for distinct receptor localization at the cell surface and distinct internalization pathways after binding DA. Finally, we targeted members of the Dop1R proximal proteome in *Drosophila* MBN *in vivo* and identified three genes that regulate R1 trafficking, as they regulate two R1 mediated behaviors – memory acquisition and starved odor attraction – and DA-mediated cAMP generation in MBNs.

## Experimental Procedures

### Cloning

The coding regions of *Drosophila* Dop1R1 (isoform D) and Dop1R2 (isoform A) were PCR amplified from pUAST-Dop1R1 (gift from Josh Dubnau) and pUAST-Dop1R2 (called pUAST-DAMB)^28^ transgenic lines, respectively, using primers with FseI restriction sites in their tail. The dresulting products were ligated into a singular FseI site in pCAGWBA, a pcDNA3.1 based expression vector^33^ to create pCAG-Dop1R1 and pCAG-Dop1R2 (untagged). The “empty” control was pCAGWBA vector with nothing inserted into the FseI site. To generate SNAP and Turbo-V5 receptor fusions, we used Infusion cloning (Takara Bio). The SNAP tag and TurboID coding sequences were PCR amplified from pSNAPf Vector (New England Biolabs, N9183S) and pTurboID-His6_pET21a plasmid (Addgene, 107177), respectively, using primers with added tails containing a flexible linker (GGSGGSGGSGGS), or not, and containing 15 bp homology to intended C or N termini of the receptors or to the pCAG vector flanking the FseI site. Tag and receptor PCR products were combined with FseI linearized pCAG and ligated together, transformed into DH5a bacterial cells prior to sequencing verification of single clones. PM-Turbo-V5 was cloned similarly but we inserted the Lyn11 peptide (MGCIKSKGKDSA)^34^ upstream of Turbo-V5 using primers and Infusion cloning.

### Transfection

We cultured and transfected HEK293T/17 cells because of high efficiency for transfection^35^, and prior characterization of *Drosophila* dopamine receptor signaling in human contexts^7^ (see Supporting information for details). Plasmids were transiently transfected using PLUS (10 ul/6 cm dish or 62.5 ul/15 cm dish) and Lipofectamine LTX (12 ul/6 cm dish or 75 ul/15 cm dish) following ThermoFisher protocol. Transfection media was added to cells for ∼21 hrs prior to characterizing expression or biotin labelling experiments. In the beginning (Figure 1A-C and S1A-E) we used equivalent DNA mass (2.52 ug/6 cm dish or 8.94 ug/15 cm dish) for empty, receptor, receptor fusion, or PM-Turbo-V5 plasmids, but found that R2-Turbo-V5 had reduced expression. For Figure S1F, as well as proteomic experiments Figure 2, we adjusted DNA mass for only R2-Turbo-V5 to be 15.75 ug /15 cm dish (PM:R1:R2, 1:1:1.76). For live-cell imaging, empty vector and dopamine receptor constructs were co-transfected with PTX-S1 as done previously^7^, along side either pGloSensor-22F (for cAMP, Promega)^36^ or CalfluxVTN (for Ca^2+^)^37^, at ratio of 6:1:6 or 6:1:1, respectively.

**Figure 1.**
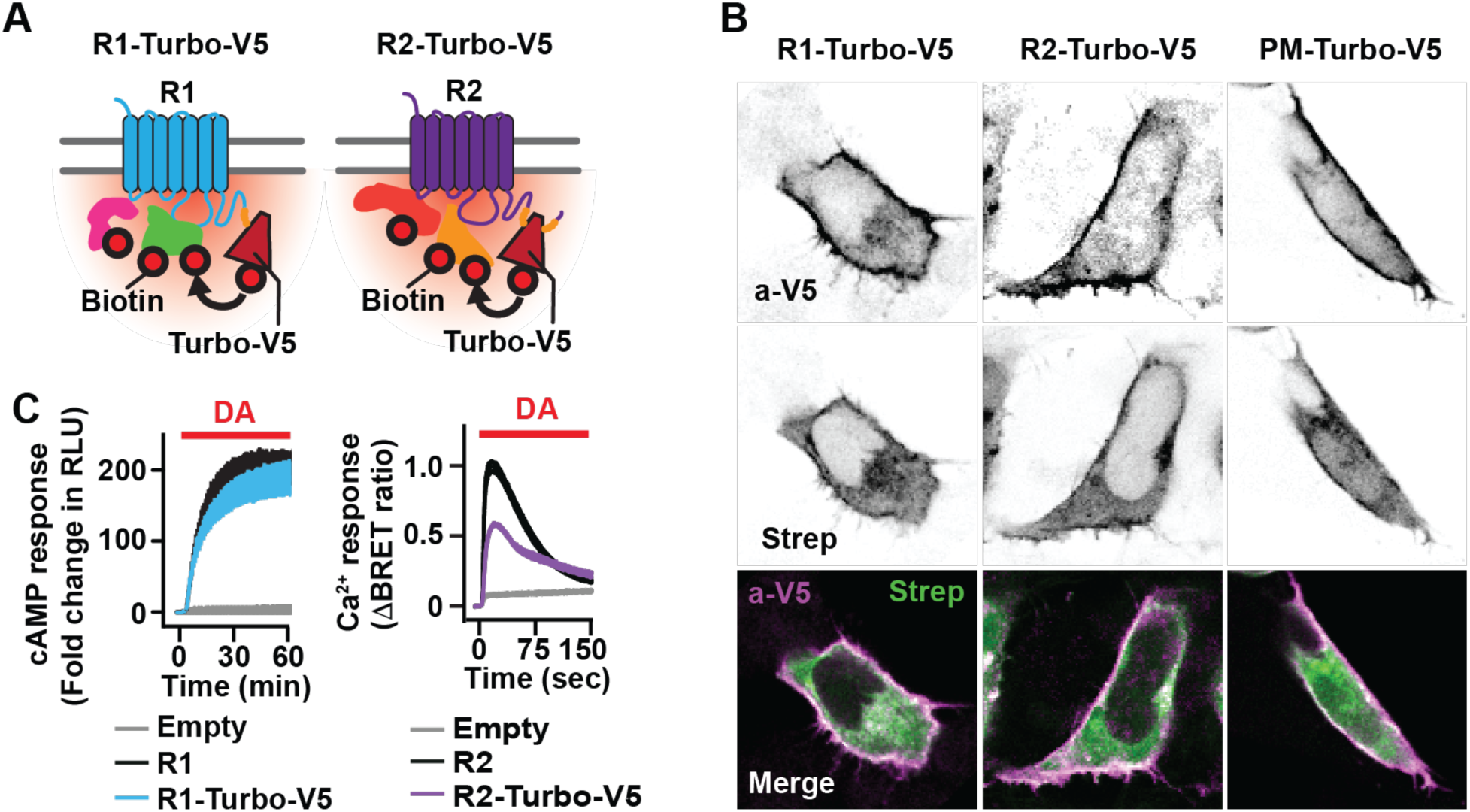
Dop1R1/R2-Turbo allows proximity labelling during receptor signaling. (A) Schematic diagram of C-terminal fusions of Turbo-V5 to Dop1R1 (R1) and Dop1R2 (R2) labeling proximal proteins with biotin. (B) Expression of R1, R2, or PM-Turbo-V5 in HEK293T cells and Turbo labelling measured by anti-V5 and Strep-488 staining, in a single Z plane. Intensity of the signal was adjusted to give similar levels across constructs for spatial comparison. (C) Live cell imaging of cAMP responses (left) and Ca^2+^ responses (right) after DA (100 μM) administration to cells expressing empty vector, R1, or R2 with or without Turbo-V5 fusion (n=6). The thickness represents the mean +/- standard error of the mean (SEM) per frame across experiments.

**Figure 2.**
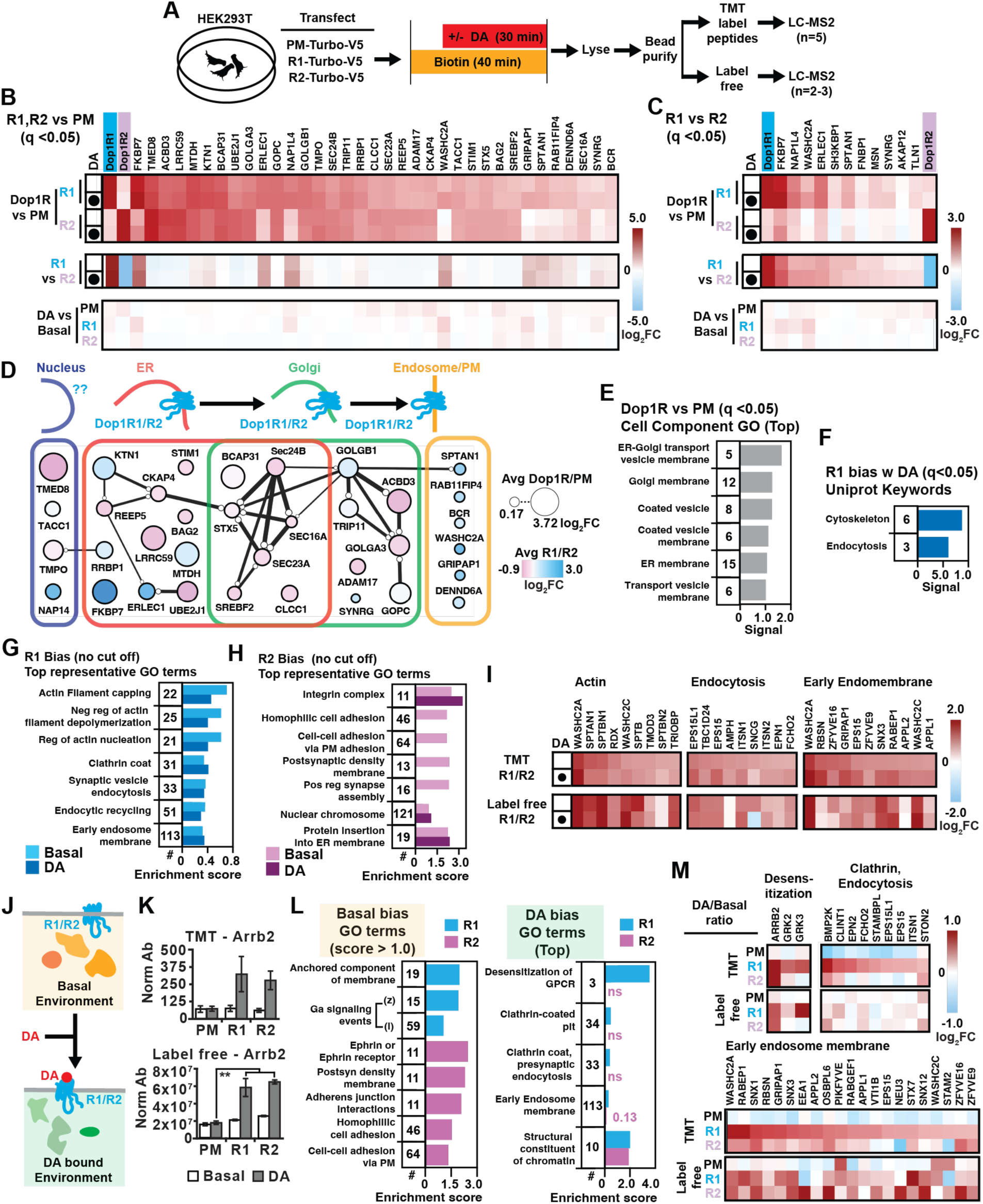
Characterization of Dop1R1/R2 proximal proteomes and ligand-mediated dynamics. (A) Schematic diagram showing cell culture-proteomics workflow used to generate TMT and label-free data sets. (B) Heatmap of abundance ratios (log2 fold change) for proteins significantly enriched (q<0.05) in R1 and R2-Turbo-V5 samples vs PM-Turbo-V5 control in the TMT data set. (C) Heatmap of abundance ratios (log2 fold change) for proteins significantly enriched (q<0.05) in R1 compared to R2-Turbo-V5 samples in the TMT data set. (D) Diagram of protein network from (B) arranged along associated cellular compartments with average abundance ratio between R1 or R2-Turbo samples vs PM-Turbo control samples (node size), the average abundance ratio between R1 and R2 samples (R1 bias blue, R2 bias purple) and annotated/potential connections (String-dg.org, minimum interaction score = 0.4, line thickness indicates strength of data support). (E) Top Cell Component GO enrichments of R1 and R2 proximal proteins from (B). (F) Uniprot Keywords GO enrichments for proteins significantly enriched in R1 vs R2 samples (R1 bias) from (C). (G,H) Top representative GO terms significantly enriched for proteins with an R1 (G) or R2 (H) bias using the String Ranked Enrichment analysis (no cut off) for Basal and DA conditions in the TMT data set. (I) Heatmap of the highest abundance ratio for R1/R2 for proteins belonging to the top representative GO terms (G) for actin, endocytosis, and early endomembrane for both TMT and label-free data sets. (J) Illustration of ligand (DA) mediated change in the proximal protein environment of Dop1R1/2. (K) Normalized abundance for Arrb2 measured across basal and DA conditions for PM, R1, and R2-Turbo-V5 samples for TMT and label-free data sets. (L) Top representative GO terms significantly enriched for proteins with a Basal (left) or DA (right) bias for both receptors using the String Ranked Enrichment analysis (no cut off) on the TMT data set. (M) Heatmap of proteins with the highest DA/Basal abundance ratio belonging to the top representative GO terms (L) for Desensitization, Clathrin, Endocytosis, and Early endosome membrane for both TMT and label-free data sets.

### Live cell imaging for cAMP and Ca^2+^

For both cAMP and Ca^2+^ signaling assays, cells were used ∼ 21 hours post transfection and reporters were imaged with CLARIOstar microplate reader (BMG Labtech) at room temperature. Details for cell processing and imaging procedures can be found in Supporting information.

### Cell staining and microscopy

For Turbo fusion and biotinylation staining (Figure S1C), 300,000 cells in 500 ul were seeded onto poly-L-lysine cover slips placed in 12 well plates and grown till ∼50-60% confluency and then transiently transfected with 1 ug/well of pCAG vectors containing Dop1R1 and Dop1R2 with or without GGS_4_-Turbo-V5 fused to the tail, as well as PM-Turbo-V5 and empty vector control. After ∼19 hrs post transfection, cells were either incubated in 500 μM biotin (using D-Biotin, Life Technologies, B20656) in normal culture media or not (normal culture media alone) for 1.5 hr in 37°C, 5% CO_2_. Biotin media was removed and cells were fixed and stained. See Supporting information for details on staining and microscopy for immuno-, biotin, and SNAP tag staining.

### Preparing Turbo labelled lysates for western blot and proteomics

For generating Turbo labelled lysates for western blot (WB) and proteomic analysis, cells were first cultured and transfected as described above using 1x 6 cm (cell #) or 4x 15 cm (cell #/dish) dishes, respectively. To measure labelling across biotin media exposure time, cells were given biotin media (500 μM, using D-Biotin, Life Technologies, B20656) for 0, 3, 10, or 40 min prior to lysation (Figure S1D) and ran in three independent experiments (quantitated in Figure S1E), 40 min exposure was chosen for the remaining WB (Figure S1F) and MS experiments (Figure 2,S2). For ligand activation specific labelling (Figure 2, S2), cells are exposed to biotin 10 min (of the 40) prior to DA exposure (via media change) to increase biotin concentrations around the Turbo environment at the time of activation. Therefore, total ligand exposure was 30 min prior to stopping the reaction and lysing the cells. After biotin and/or DA treatment, cells were processed to make lysates for bead incubation or western blot analysis (see Supporting information).

### Streptavidin bead purification

Bead purification based on Turbo related methods in Branon et al., 2018. Briefly, streptavidin magnetic beads (Dynabeads MyOne Streptavidin T1, ThermoFisher, #65601) were pre-washed as recommended by the manufacturer using the above lysis buffer, and then incubated with lysates in a microfuge tube (1 μl beads/ 4.3 μg protein) on a nutator on random agitation with end over end rotation for 16-24 hr at 4°C. Beads were then washed and separated at 4°C in the following solution/sequence: lysis buffer (x2), 1M KCl (x1), 100mM Na_2_CO_3_ (x1), 2M Urea in 10mM Tris-HCl pH 8 (x1), and lysis buffer (x2). Proteins were then eluted from the beads by resuspension in elution buffer (0.1% SDS, 2mM biotin) and then beads were vortexed for 10 sec, boiled at 95°C for 3 min, repeated 3 times. After separation, the eluants contained ∼20-40 ug purified biotinylated proteome per sample measured by BCA assay.

### LC-MS/MS protein identification and quantification

For the TMT data set, eluted proteins (6 experimental conditions, 5 biological replicates/condition) were precipitated overnight with 4X ice cold acetone (v:v). Protein pellets were then processed for trypsin digestion using micro S-TrapsTM (Protifi, Huntington, NY) according to the manufacturer’s instructions. Briefly, the dried protein was resuspended in 22 μL of 5% SDS, reduced using DTT at 55°C for 10 minutes, alkylated using methyl methanethiosulfonate (MMTS) at room temperature for 10 minutes, and then spun down at 20K for 10 minutes. Subsequently, 2.5 μL of phosphoric acid was added to the sample, followed by 165 μL of a mixture of HPLC grade methanol and Protifi binding/wash buffer. Following loading onto the S-TrapTM, samples were washed three times using centrifugation and trypsin was added in 50 mM TEAB at a 1:25 w/w ratio. The S-TrapTM column was incubated for 1 hour at 47°C. Following this incubation, 40 μL of 50 mM triethylammonium bicarbonate (TEAB) was added to the S-TrapTM and the peptides were eluted using centrifugation. Elution was repeated once. A third elution using 35 μL of 50% acetonitrile (ACN) was also performed and the eluted peptides were dried under vacuum. Peptides were reconstituted in 50 mM TEAB and their concentrations were determined using the PierceTM quantitative fluorometric peptide assay (Thermo Fisher Scientific, Waltham, MA). Six micrograms of peptides were labelled with TMT labels (6-plex) according to the manufacturer’s instructions (Thermo Fisher Scientific, Waltham, MA) and pooled. The pooled plexed samples were dried under vacuum, resolubilized in 1% TFA and desalted using OASIS HLB 1cc solid phase extraction cartridges (Waters, Milldford, MA), and then dried once again using vacuum. The dried labelled peptide plexed samples were then resuspended in 400 μL 1M TEAB and fractionated using an Agilent 1100 HPLC system and high pH reversed phase chromatography on a Zorbax Eclipse XDB-C18 column (4.6 x 150mm, 5-micron) from Agilent (Santa Clara, CA). Sixty fractions were collected at 0.5 mL/min over 105 min using a gradient of 0-5% solvent B in 10 min, 5-35% solvent B in 60 min, 35-70% solvent B in 15 min, a 10 min hold of 70% solvent B, and finally a return to 5% solvent B in 10 min. Solvent A consisted of 100mM TEAB and solvent B consisted of ACN. The 60 fractions were dried vacuum, concatenated into 6 fractions, and then cleaned up using 2 μg capacity C18 ZipTips (Millipore, Billerica, MA) according to the manufacturer’s instructions and dried.

Dried TMT-labelled peptides were reconstituted in 5 μL of 0.1% formic acid and on-line eluted into a Fusion Tribrid mass spectrometer (Thermo Scientific, San Jose, CA) from an EASY PepMapTM RSLC C18 column (2μm, 100Å, 75 μm x 50 cm, Thermo Scientific, San Jose, CA) using a gradient of 5-25% solvent B (80/20 acetonitrile/water, 0.1% formic acid) in 180 min, followed by 25-44% solvent B in 60 min, 44-80% solvent B in 0.1 min, a 5 min hold of 80% solvent B, a return to 5% solvent B in 0.1 min, and finally a 20 min hold of solvent B. All flow rates were 250 nL/min delivered using a nEasy-LC1000 nano liquid chromatography system (Thermo Scientific, San Jose, CA). Solvent A consisted of water and 0.1% formic acid. Ions were created at 1.7-2.4 kV using an EASY Spray source (Thermo Scientific, San Jose, CA) held at 50oC. A synchronous precursor selection (SPS)-MS3 mass spectrometry method was used by scanning between 380-2000 m/z at a resolution of 120,000 for MS1 in the Orbitrap mass analyzer, and performing CID at top speed in the linear ion trap of peptide monoisotopic ions with charge 2-8, using a quadrupole isolation of 0.7 m/z and a CID energy of 35%. The top 10 MS2 ions in the ion trap between 400-1200 m/z were then chosen for HCD at 65% energy and detection in the Orbitrap at a resolution of 60,000 and an AGC target of 1E5 and an injection time of 120 ms (MS3).

Quantitative analysis of the TMT experiments was performed simultaneously to protein identification using Proteome Discoverer 2.4 software. The precursor and fragment ion mass tolerances were set to 10 ppm, 0.6 Da, respectively. Enzyme cleavage was performed by Trypsin with a maximum of 2 missed cleavages and Uniprot Human proteome FASTA file (downloaded in 2017) with six additions (P00761|TRYP_PIG, P22629|SAV_STRAV, P02769|ALBU_BOVIN, P06709|BIRA_ECOLI, Q24563|DOPR2_DROM, P41596|DOPR1_DROME) was used in SEQUEST searches. The impurity correction factors obtained from Thermo Fisher Scientific for each kit was included in the search and quantification. The following settings were used to search the data; dynamic modifications; Oxidation / +15.995Da (M), Deamidated / +0.984 Da (N, Q), Biotin / +226.078 Da (K), TMT6plex / +229.163 Da (K) and static modifications of TMT6plex / +229.163 Da (N-Terminus), Carbamidomethyl +57.021 (C). Only unique+ Razor peptides were considered for quantification purposes. Percolator feature of Proteome Discoverer 2.4 was used to set a false discovery rate (FDR) of 0.01. Co-isolation threshold and SPS Mass Matches threshold were set to 50 and 65, respectively. Total Peptide Amount normalization method was used to correct for loading bias. S/N (signal to noise ratio) processed through Protein Abundance Based method to calculate the protein level ratios. The Label free data set was generated as described in the Supporting information.

### Gene ontology analysis

First, proteins that were significantly more abundant around either R1 or R2 comparing receptor-Turbo samples to PM-Turbo-V5 (based on abundance ratios, q <0.05) were ran on String-db.org database as “Multiple proteins” using their Uniprot accession numbers and statistically significant functional enrichments were identified (Figures 2E,F; Supporting spreadsheet 2). Next, a Ranked functional enrichment analysis (“Proteins with Values/Ranks”, String-db.org) was performed comparing proteomes consisting of all proteins detected and quantitated in at least 2 biological replicates (6164 proteins across all TMT samples, 5779 proteins for all label free samples). For each comparison, the proteomes were ranked according to their mean raw abundance ratios (e.g., R1^Basal^ / PM^Basal^) and a high (1 percent) FDR stringency was used to identify functional gene pathways or gene ontologies that are statistically enriched taking into account the number of members present and their position toward the end of the raw abundance ratio distribution across the sample proteome (Figures 2G,H,L; Supporting spreadsheet 2).

### Fly husbandry

Flies were maintained on standard fly food at 25°C. UAS-RNAi stocks were obtained from Vienna Drosophila Stock Center (http://stockcenter.vdrc.at/control/main) and Bloomington Drosophila Stock Center (https://bdsc.indiana.edu/). All RNAi lines used in this study are listed in Supplemental Table 1). We employed the control lines used to develop the UAS-RNAi transgenic line in all experiments: KK60100, GD60000, TRIP36303, and TRIP36304. For all RNAi experiments, lines were crossed to; UAS-Dicer2; R13F02-Gal4. For behavioral experiments, a mix of males and females were used. For physiological experiments, MB-^T^epac^VV^ cAMP reporter or UAS-GRAB(ACh) sensor was used and only females were imaged.

### RNAi screen selection

The human Dop1R proximal proteome (Figure 2B) and Arrb2 (41 proteins) were searched through DIOPT (https://www.flyrnai.org/diopt)^39^ for predicted orthologous *Drosophila* proteins based on sequence. We chose *Drosophila* orthologs that had a) moderate to high “homology” b) measurable expression in any subclass of MBN^40^ and c) “good” quality RNAi (targeting coding or 3’UTR sequence, all isoforms, and no predicted off target effects). The RNAi screen consisted of 48 RNAi targeting 48 genes due to more than one *Drosophila* gene with moderate to high homology for a few human genes.

### Aversive Olfactory Paradigm

Flies 1-4 day old were used for standard olfactory aversive conditioning experiments. Aversive olfactory behavioral experiments were conducted under red light, and in ∼65–75% humidity at 23–25°C. Flies were tapped into new food vials and acclimated for 15 minutes to behavioral testing conditions before training. Groups of ∼70 flies were tapped into tubes where they received 30 seconds of air, 1 minute of CS+ odor paired with electric shock (12 shocks 1.25sec each, at 5sec intervals at 90V), 30 seconds of air, 1 minute of CS- odor with no electric shock, and 30 seconds of air. After the specified time (3min or 3hr), flies were loaded into a T-maze and given 1 minute to acclimate, then given 2 minutes to choose between the arms containing CS+ odor and CS- odor^41^. The number of flies in each arm were counted and used to calculate a preference index ((CS-) – (CS+))/((CS-) + (CS+)). Odors used were counterbalanced and included 3-octanol (OCT; 0.05%) and 4-methylcyclohexanol (MCH; 0.07–0.1%) diluted in mineral oil. Odor concentrations were adjusted slightly each day so that flies displayed similar aversion scores to both odors when tested in the T-mazes.

### Innate Odor Avoidance Paradigm

Flies 0-3 days old were collected. Fed flies were maintained on normal fly food until testing. Starved flies were moved to vials with only a Kimwipe soaked with 2 mL of deionized water 48–50 hours prior to testing. Flies were transferred to a T-maze and given 1 minute to acclimate, then given 2 minutes to choose between the 2 arms, one delivering MCH (0.08%) diluted in mineral oil and the other delivering mineral oil only. A performance index was calculated for each group using the formula (number of flies in the odor arm) - (number of flies in non-odor arm)/total flies in both arms.

### Fixed brain immunostaining

The brains of 4-6 day old females were dissected in cold S2 media and immediately placed in 1% paraformaldehyde in S2. Brains were fixed overnight in paraformaldehyde and then washed PAT3 (0.5% BSA, 1X PBS, 0.5% Triton X-100) for 30 minutes at a time. Blocking was performed by washing brains with PAT3 + NGS (normal goat serum) for 1 hour before primary antibodies. Brains were incubated in primary antibodies (rabbit anti-GFP 1:1000, mouse anti-NC82 1:50) for 3 hours at room temperature and overnight at 4 degrees and secondary antibodies (488 Alexa anti-rabbit 1:1000 and 633 Alexa anti-mouse 1:1000) for the same duration. Brains were mounted in Vectashield and imaged using 488nm laser on Zeiss Axio Observer Z1 inverted confocal microscope. Images were analyzed using imagej.

### *Ex vivo* Functional Imaging

The brains of 4–6-day old females were mounted flat in a recording chamber, such that the anterior face was facing the microscope objective^42^. Time series images were collected over a period of 8 minutes at 0.5 Hz. Brains were treated with saline containing 1µM tetrodotoxin (TTX) 10 minutes before recording started. Dopamine was applied by switching the solution perfusing the brain from saline to 10 mM dopamine dissolved in saline for 30s. The levels of cAMP in response to stimulus were reported in terms of the normalized change in inverse FRET ratio (ΔR/R0 = (R-R0)/R0 where R = CFP emission / YFP emission). R0 represent the baseline fluorescence within the ROI. Data were analyzed from stable recordings with no major visible drift using ImageJ software.

### Statistical analysis

For TMT proteomics data, an ANOVA was used to identify proteins that are regulated across comparison groups (R1 or R2 vs PM, Basal vs. DA for each receptor, and R1 vs. R2 for Basal and DA). The multiple testing correction as per Benjamini Hochberg (B-H) was applied to identify a “top tier” of significant proteins and limit identification of false positives with a False Discovery Rate (FDR) of 5 percent. Adjusted p values, or q-values, of <0.05 were considered statistically significant. For behavioral data, learning and avoidance indices were calculated as described above in “aversive olfactory paradigm” and “innate odor avoidance paradigm” respectively. One-way ANOVA with a Bonferroni post hoc analysis was performed to compare performance indices. For ex vivo cAMP responses to DA, FRET ratios were calculated as described above in “ex vivo functional imaging”. Mann-Whitney U test with Bonferroni correction for multiple comparisons was used. All tests were two tailed and confidence levels were set at α = 0.05.

## Results

### Proximity labelling of Dop1R1 and Dop1R2 in cells using tail Turbo fusion

We first chose to use the efficient and live cell capable TurboID proximity labelling system^16^ by fusing the Turbo ligase to the *Drosophila* D1-like DA receptors and express them in HEK293T cells. This approach takes advantage of the conserved signaling for R1 and R2 between human cells and MBN synapses^7,8^ to identify candidate memory regulators occupying receptor proximal environments in basal- and ligand-driven signaling and transport. We reasoned that we could identify conserved proteins that we could interrogate *in vivo* in *Drosophila* memory circuits.

First, we tested the optimal location for making fusions to R1 and R2 by evaluating the resulting DA-mediated signaling downstream of these receptors using live-cell imaging. We created human cell line expression plasmids (pCAG) containing Dop1R1 (isoform E) and Dop1R2 (isoform A) open reading frames, and we additionally inserted a small chemical tag SNAP (New England Biolabs)^43^ with or without a linker (GGS x4) at either the N-terminus or in the cytoplasmic tail of the receptors. To test the resulting impact on receptor function, we utilized live cell optical imaging of cAMP (pGLoSensor, Promega)^36^ and intracellular Ca^2+^ (CalfluxVTN)^37^ via transient co-transfection of reporter plasmids with R1 (or R2)-SNAP plasmids in HEK293T cells, and applied DA (100μM)^7^. For both receptors, N-terminal localization of the SNAP tag, regardless of the presence of a linker, led to a large impairment of DA-mediated cAMP (top) for R1 and R2, and Ca^2+^(bottom) for R2 (Supplemental Figure 1A). Notably, tagging the tail with SNAP tag and a GGS^4^ linker had minimal impact on R1 and R2 signaling (cAMP or Ca^2+^), and the tagged receptors were expressed at the plasma membrane (Supplemental Figure 1B). Therefore, we found that fusions of a tag to the C-terminal tail including a flexible linker best maintained the R1/R2 function.

We next tagged the C-terminus of R1 or R2 with the intended, but larger, Turbo (+V5 peptide) sequence (Figure 1A). As a control for ubiquitous Turbo labelling at the plasma membrane and non-specific Turbo-V5 peptide interactions, we also made a receptor-less Turbo control that included Turbo-V5 targeted to the plasma membrane (PM-Turbo-V5) via Lyn kinase N-terminal peptide^34^. Transient transfection of R1 and R2-Turbo-V5 constructs led to robust expression (via anti-V5 signal) along the plasma membrane, at outstretched protrusions, and internally, in the endomembrane system, compared to empty vector transfection (Figure 1B, S1C). PM-Turbo-V5 transfection also led to a preference for V5 staining along the plasma membrane and protrusions, similar to R1 and R2-Turbo-V5 expression. Biotin media (500 μM) exposure for 1.5 hr led to a robust biotinylation of cellular proteins (via Strep-AF488 dye), with overlap particularly at the plasma membrane. Biotin staining internally with no Turbo-V5 overlap is expected due to diffusion after labelling. Furthermore, Turbo-V5 fusion did not disrupt R1-mediated DA-> cAMP signaling, but, unlike the smaller SNAP tag, it did reduce DA- >R2->Ca^2+^ signaling by ∼50% (Figure 1C). We noted that comparing V5 staining and biotin-labelled proteomes created by transfection with equal mass of pCAG plasmids containing PM, R1, or R2-Turbo, resulted in similar levels of PM and R1-Turbo-V5 expression, but R2-Turbo-V5 levels and resulting Strep labelling were markedly reduced (Figure S1C). It is possible that Turbo fusion to R2 reduced the stability of its mRNA or protein and this reduced the R2 live-cell signaling we observed in Figure 1C. To account for this in our proteomics experiment, we adjusted the amount of DNA during transfection for PM, R1, or R2-Turbo-V5 to obtain similar amounts of biotinylated proteome after 40 min Biotin exposure (Figure S1D).

As has been reported^16^, we found that Turbo-mediated biotinylation occurs without biotin supplementation, likely originating from biotin in the standard culture media, but biotin supplementation (500μM, 1.5 hr) robustly increases labelling (Figure 1B, S1C). Using a shorter time course, we found that biotin addition increased labelling after 10 min and increased further until ∼40 min (Figure S1E,F). Overall, our system allows Turbo labelling of Dop1R environments and can be used to characterize active DA-mediated signaling.

### Dop1R proximal proteomes span receptor trafficking compartments

To characterize the basal and ligand driven protein networks and environments around Dop1R1/2 receptors, we expressed R1-Turbo-V5, R2-TurboV5, or the PM-TurboV5 control, then applied biotin media (500μM) for 40 min (Figure 2A). During the 40 min of biotin, we allowed 10 min for biotin to penetrate the receptor environments, a timepoint Turbo labelling begins (Figure S1E,F), before applying DA for 30 min (“DA”) or not (“Basal”). The cells were lysed and biotinylated proteomes were bead-purified and digested into peptides. These peptides were processed in one of two methods prior to LC-MS2 (See Methods). For our primary data set, the peptides were labelled with an independent TMT tag per condition (6 tags / 6 conditions), and all conditions were then combined for 5 replicates prior to fractionation and LC-MS2. As an independent approach, we repeated the experiment with label-free quantitative proteomics that involved no fractionation before LC-MS for all 6 conditions for 2-3 replicates (1 sample was lost). A total of 6164 proteins and 5779 were quantitated across all conditions and replicates in our TMT and label-free data sets, respectively (Figure S2A, Supporting spreadsheet 1). Despite fundamentally different post-processing of the peptides and fractionation or not, ∼78% of the TMT proteome is detected in the label-free data set. Furthermore, abundance ratios (abs(log2 ratio) > 0.5) between conditions (e.g., R1 Basal/PM Basal) for TMT proteins were positively and linearly correlated with their abundance ratio in the label-free data (Figure S2C), indicating remarkable reproducibility of data sets. However, given the increased quantitative power of TMT proteomics and its larger sample size, going forward, we use the label-free data as support for findings from the more statistically robust TMT data set.

Using standard multi-test correction q-value analysis on our TMT data set, we found 35 human proteins were significantly more abundant in R1-Turbo or R2-Turbo samples (basal and DA) compared to the receptor-less plasma membrane bound PM-Turbo control (q<0.05, Figure 2B), and 11 human proteins whose R1/R2 abundance ratio was significantly different, all of which are biased to be labelled by R1-Turbo (Figure 2C). Many of the 11 R1 specific proteins are also significantly more abundant in R1 expressing samples compared to PM control (Figure 2B). Importantly, the *Drosophila* proteins Dop1R1 and Dop1R2 themselves were robustly present only in conditions where the respective receptor was expressed and similar levels between basal and DA conditions (Figure S2D), indicating Dop1R-Turbo self-labeling, validating our Turbo methodology. Remarkably, most of the TMT significant proximal proteins also have similar abundance ratios between TMT and Label-free data sets as seen on volcano plots (Figure S2E, F). Gene ontology (GO) analysis (String.db) on this significant Dop1R proximal network shows significant enrichment for proteins found in cellular locations expected along the transport/lifespan of these GPCRs, including the ER, Golgi, Endosome, and PM (Figure 2D,E, Supporting spreadsheet 2). This further validates that we indeed captured Dop1R proxiomes and potential interactors.

### R1 and R2-Turbo are localized to distinct subcellular environments

R1 and R2 drive distinct cellular signaling in human cells and synapses^7,8^, indicating that the receptors are located in differential signaling environments and subcellular locations. We first compared R1 and R2-Turbo proteomes using GO analysis with the small 11 protein network that is significantly (q<0.05) more abundant around R1 than R2. Given the small network, we did not find significant enrichments in traditional GO categories, but we did find two significant enrichments for Uniprot Keywords ‘Cytoskeleton’ and ‘Endocytosis’, suggesting that R1 is proximal to pathways involving these keywords (Figure 2F). To fully utilize our two data sets and gain more insights into differences between R1 and R2 function, we performed a Ranked Functional Enrichment analysis (String-db.org) using all quantified proteins (no cutoffs) ranked by raw abundance ratios (6164 proteins for TMT, 5779 for Label-free). This analysis utilizes the entire dataset to find significantly enriched pathways based on the number and abundance ratio values of pathway members with respect to the ends of the whole distribution, and we used our data sets ranked by R1/R2 ratios (Figure 2G, H, Supporting spreadsheet 2). We found that the top significant enrichments with an R1 bias included actin regulation, clathrin, synaptic vesicle endocytosis, or were located at the early endosome membrane for both data sets (Figure 2G, Supporting spreadsheet 2). This corroborates our 11-protein q-value analysis on the cytoskeleton and endocytosis pathways (Figure 2F). Many proteins in these pathways have positive R1/R2 abundance ratios (Top R1/R2 basal ratio proteins, Figure 2I) and large average normalized abundance around R1-Turbo compared to R2-Turbo and PM-Turbo (Figure S2G). This pattern is remarkably conserved across TMT and label-free data. In particular, clathrin based endocytosis regulating proteins FNBP1 and SYNRG are significantly more abundant in R1 than R2 samples (Figure 2C) as well as EPS15 and EPS15L1 (not significant) have large R1/R2 ratios in both data sets (Figure 2I, S2G). Furthermore, the WASHC2A protein involved in endosomal sorting and recycling back to the plasma membrane is significantly more abundant in R1 samples than R2 (Figure 2C,I, S2G), while WASHC2C did not reach q value significance but was clearly enriched in R1 vs R2 (Figure 2I, S2G). This indicates R1 might be sorted and recycled in the endosomal system differentially than R2. Overall, R1 appears to be regulated at the membrane via canonical clathrin-mediated synaptic vesicle endocytosis, and R1’s proximity to cytoskeleton regulatory proteins, at least in part, reflects the importance of the cytoskeleton in vesicle endocytosis and sorting in the endosome^44^.

Despite R2-Turbo localization at the plasma membrane (Figure 1B, S1B, C), proteins related to synaptic vesicle and clathrin-mediated endocytosis were labelled far less by R2-Turbo (Figure 2I, S2G), suggesting R2 might be internalized differentially from R1. Interestingly, the top significant enrichments with an R2 bias included integrins, cell adhesion, and post-synaptic density environments for both data sets (Figure 2H, S2H, Supporting spreadsheet 2). We noted that the top 2 post-synaptic density proteins with R2 bias were Ephrin type-B receptor 2 (Ephb2) and neuroplastin (Nptn) (Figure S2H). Both Ephrin signaling and Nptn mediate cell-cell signaling that play a role in synaptic plasticity underlying memory in mammals and *Drosophila*^45–48^. We also observed bias towards labelling nuclear chromosome-associated proteins, like histone H2AX, with R2-Turbo compared to R1-Turbo (Figure 2H; S2H), however, this was not reproduced in the label-free data set. Whether this indicates R2’s localization near the nucleus or nuclear proteins’ localization near the plasma membrane remains unclear. Overall, these results indicate R1 and R2-Turbo are localized in separate subcellular compartments near or at the plasma membrane, which could support their distinct cellular functions in synapses.

### DA differentially shifts R1 and R2 signaling environments

Ligand binding to a GPCR usually drives its subsequent internalization (or endocytosis) away from the plasma membrane, and this desensitizes the cells response to subsequent ligand^49,50^, and this has been seen to occur in minutes and proximity labelling has been used to observe this movement for other receptors^18^. Desensitization involves phosphorylation by G-protein receptor kinases (GRKs), binding to B-Arrestin2, internalization, and movement of the receptor to the early endosome^51,52^. Thus, we aimed to use our Dop1R-Turbo system to capture the movement of these receptors from “Basal” environments to DA-bound environments (Figure 2J). Comparing ligand (DA) driven and basal proteomes within the TMT data set revealed no protein reached significance with our strict q<0.05 cutoff for any Turbo condition. However, interestingly, Beta-Arrestin2 (Arrb2) was robustly enriched (∼2 fold) when either dopamine receptor *and* DA were simultaneously present, but low in all other conditions (Figure 2K, top). In the label-free data set, Arrb2 was similarly enriched with DA+Dop1R samples and was significantly increased for both receptors in the presence of DA compared to PM-Turbo + DA (Figure 2K, bottom).

Given Arrb2’s role in ligand-driven endocytosis of GPCRs, these results are encouraging that we captured DA-driven interactions, and thus we proceeded to run a Ranked Functional Enrichment analysis on all quantitated proteins ranked by raw abundance ratio (DA/basal). Proteins with high DA/basal ratios should be enriched for new signaling environments driven by ligand activation (DA bias), and those with low DA/basal ratios would be enriched for old signaling environments left behind after ligand activation (Basal bias). We found that many GO terms were significantly enriched with either Basal or DA bias and were both common and unique for both receptors for the TMT data set (Figure 2L; Supporting spreadsheet 2). Importantly, we found that for R1-Turbo, the top significantly enriched pathways with a Basal bias included anchored component of the membrane, GPI-anchoring, and Gaz and Gai signaling events (Figure 2L, left). Many proteins in the TMT data set associated with these pathways had reduced abundance in the presence of DA for R1-Turbo, and to a lesser extent, the label-free data set (Figure S2I). In particular, SMPDL3B, PRNP, CD44, CD55, proteins targeted to lipid rafts via GPI-anchor, were markedly reduced in R1 samples when DA is present (Figure S2I), suggesting R1 resided in lipid raft microdomains in the plasma membrane before being internalized after activation. Regarding R2, we found in the TMT data set the top significantly enriched pathways with a Basal bias included similar pathways as those with R2 bias (Figure 2H,I), specifically ephrin signaling, post-synaptic density, and cell adhesion pathways (Figure 2L). Therefore, unlike R1, DA drives R2 away from cell-cell adhesion-like signaling environments at the membrane.

On the other hand, the top significantly enriched pathways for R1 with a DA bias in the TMT data set included GPCR desensitization (Figure 2L, right) that includes only ARRB2, GRK2, and GRK3, which in the case of R1 had increased abundance in the presence of DA, whereas for R2 only ARRB2 increased its abundance with DA, and this pattern was remarkably consistent across both data sets (Figure 2M, S2J). Thus, R1 and R2 have differential engagement with GRKs when activated by DA. Additionally, the pathways of clathrin-coated pit, presynaptic endocytosis (Figure 2L, right), and early endosome membrane (Supporting spreadsheet 2) were significantly enriched when DA is present for R1 (but not R2) with many of proteins having increased their abundance in the presence of DA in R1 samples (Figure S2J). Altogether, these data indicate that Turbo labelling captured DA-mediated shifts in R1’s location away from the plasma membrane and towards clathrin mediated internalization pathways and locations.

Looking at the DA bias regarding R2, we found, similar to R1, a significant enrichment for the early endosome membrane with a DA bias. Therefore, this result, combined with our findings that DA drives R2’s proximity/interaction with Arrb2 (Figure 2K), indicates that DA drives R2 internalization away from the membrane to the early endosome, albeit not through clathrin-mediated endocytosis like R1.

Finally, we found that in the TMT data set, both R1 and R2 show significant enrichment for chromatin-related proteins with a DA bias, and these consist mostly of histones (Figure 2L, S2K), like nuclear chromosome proteins having R2 bias above (Figure S2H). This was again unexpected and could indicate that histones are moonlighting outside of the nucleus near the plasma membrane, or ligand activation of these dopamine receptors drives localization inside the nucleus^53,54^. However, we did not replicate this DA-mediated increase in abundance of these chromatin proteins in the label-free data set. Nevertheless, our results indicate that our proximity labelling approach was successful in capturing ligand-driven shifts in receptor location and signaling, suggesting fundamental differences between these receptors in their plasma membrane localization (GPI anchor vs cell adhesion) in basal conditions, DA driven engagement with GRKs, and endocytosis pathway (clathrin or not) when activated by DA.

### Trafficking proteins identified to play a role in two dopamine-mediated behaviors

Given that R1 was proximal to cell components and pathways related to vesicle and GPCR trafficking for both data sets (Figure 2I, S2, Supporting spreadsheet 2), we sought to identify R1 trafficking proteins that regulate DA-mediated learning in *Drosophila*. First, we targeted the *Drosophila* orthologs of the human Dop1R proximal network (39 proteins, q<0.05, Figure 2B) and Arrb2 in the MBN using R13F02-Gal4 driven Dicer2 and RNAi expression (Figure 3A), and we tested R1-dependent aversive odor learning (Figure 3B). During training, animals were first exposed to one of two odors (3-Octanol, OCT, or Methylcyclohexanol, MCH) simultaneously with 6 electric shocks (moderate training), followed by exposure to the other odor. To test learning, we tested memory immediately after training. Disruption of many orthologs of Dop1R proximal proteins tended to enhance or disrupt immediate memory (Figure 3C), as their mean +/- SE did not overlap with the mean +/- SE of the same day control (#). Among these, we noted disruption of two proteins related to ER->Golgi transport, Sec24AB and CG13887, which disrupted and enhanced memory, respectively. Furthermore, disruption of krz, an ortholog of human Arrb2 and likely a regulator of endocytosis, tended to reduce learning^55^. Similar to our proteomics results for Arrb2 (Figure S2K), human orthologs for Sec24AB (human Sec24B) and CG13887 (human BCAP31) showed significant enrichment in R1 and R2-Turbo-V5 samples regardless of DA application in the TMT data set (and also for BCAP31 in the label-free data set as well)(Figure S3A).

**Figure 3.**
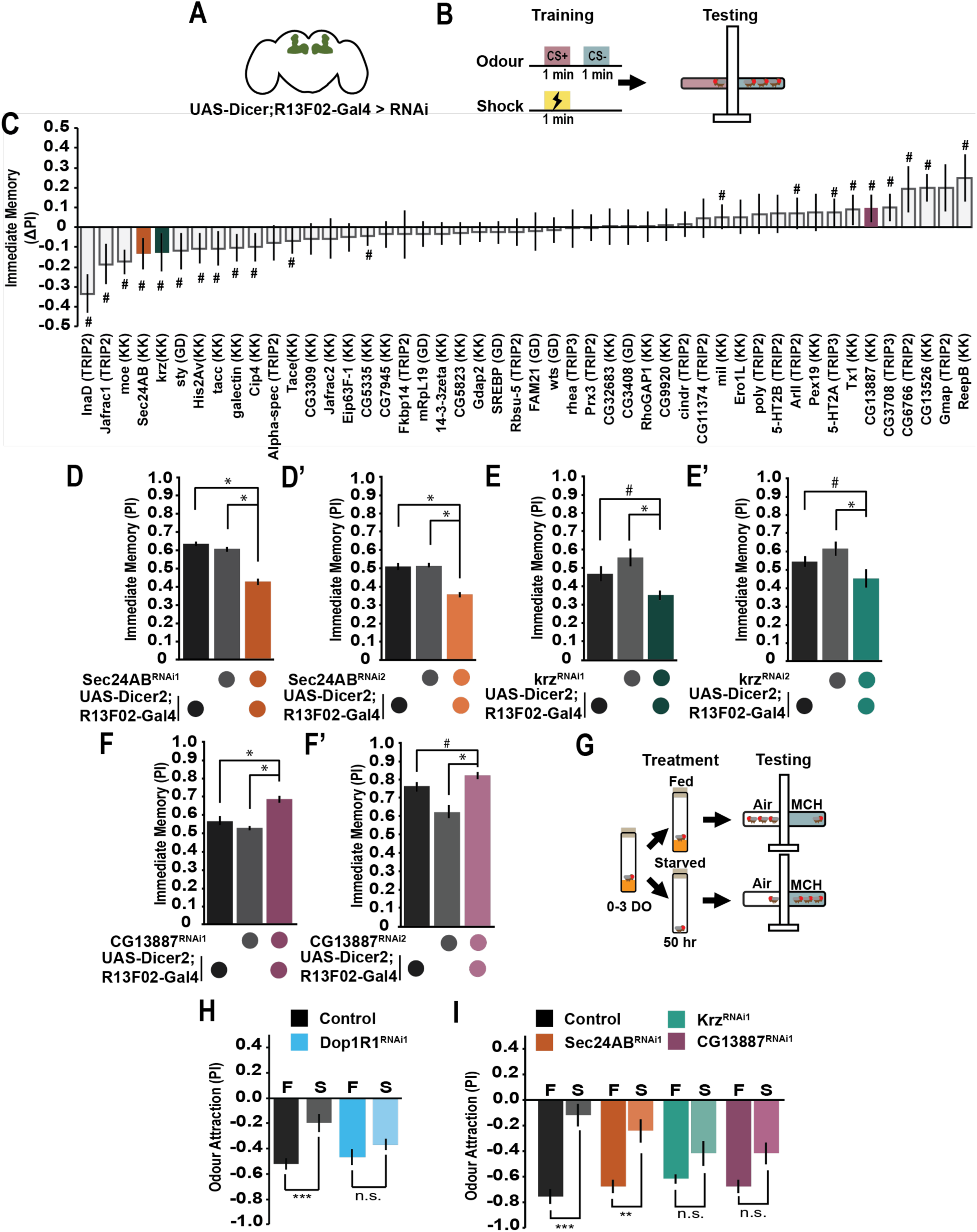
Trafficking proteins identified to participate in dopamine-mediated learned and state-dependent innate behaviors. (A) Image of the pan-MBn driver (R13F02-gal4) used for the learning and memory screen. Cartoon representation of R13F02 expression. (B) Schematic of the aversive olfactory paradigm used for the learning and memory screen. (C) Small-scale RNAi screen performed on 48 proteins discovered in the proximity labelling experiment. Learning was tested, and a delta performance index (PI) was calculated by subtracting the PI of the experimental line from the PI of an internal control tested at the same time as the experimental line, with an N=4. An # symbol was used to represent RNAi lines whose error bars did not overlap with the error bars of the control. (D-D’) Sec24AB knockdown in the MBn decreased immediate learning. Controls: R13F02>+ and Sec24AB^RNAi^>+. (N=8, one-way ANOVA, Tukey post hoc, * = p<0.05, mean ±SEM). (E-E’) krz knockdown in the MBn decreased immediate learning. Controls: R13F02>+ and Krz^RNAi^>+. (N=8, one-way ANOVA Tukey post hoc, * = p<0.05, # = p<0.1, mean ±SEM). (F-F’) CG13887 knockdown in the MBn enhanced immediate learning. Controls: R13F02>+ and CG13887^RNAi^>+. (N=8, one-way ANOVA Tukey post hoc, * = p<0.05, # = p<0.1, mean ±SEM). (G) Schematic of the starved odor preference paradigm to test innate odor avoidance behaviors. (H) Expression of Dop1R1 RNAi in the MBn significantly altered the starved odor attraction to MCH compared to starved control flies (N=8, Student’s t-test, * = P<0.05). (I) Fly odor avoidance was tested to MCH in both a fed and starved state. RNAi was expressed in the MBns using R13F02-gal4 (N=8, students t-test, * = P<0.05, **=P<0.01, *** = P<0.001).

To validate these potential R1 trafficking proteins’ role in learning, we targeted Sec24AB, CG13887, and Krz in the MBN using the screen RNAi line and an additional RNAi line targeting an independent portion of each gene. Disruption of Sec24AB and Krz in the MBN significantly decreased immediate memory with both RNAi lines compared to two genetic controls (1 lacking Gal4, 1 lacking RNAi)(Figure 3D, E). Remarkably, disruption of CG13887 in the MBN significantly enhanced immediate memory for both RNAi lines compared to controls (Figure 3F, F’). Finally, disruption of these genes in the MBN did not impair odor or shock avoidance behavior, with the exception that odor avoidance to only one odor (MCH) was slightly but significantly reduced for Krz (Figure S3B, C).

To evaluate how these genes regulate the acquisition of associative memory across training intensity, we varied the number of electric shock punishments coinciding with the odor. We found that disruption of Sec24AB in the MBN did not disrupt immediate memory at the lowest training intensity (1 electric shock), but did significantly decrease immediate memory with 2 electric shocks and above (Figure S3D, top), suggesting Sec24AB is required for strengthening the association with additional punishments, but not required for the initial association. Disruption of Krz in the MBN significantly decreased only the highest intensity training (Figure S3D, middle), indicating that it is required for maximal memory strength. Finally, disruption of CG13887 enhanced immediate memory at 6 and 12 electric shock associations, suggesting CG13887 normally limits the maximal strength of the association and is a memory suppressor gene^56^.

Krz is one of two non-visual Arrestins in *Drosophila*, alongside CG32683. We targeted CG32683 in the MBN and did not see a significant decrease in learning (Figure S3E). This suggests Krz might be the primary non-visual Arrestin supporting learning. Sec24AB, and its paralog Sec24CD, in flies are critical components of the COPII pathway and predicted to be two adaptor proteins that recognize distinct GPCR cargo for ER export, alongside cargo invariant COPII components like Sec16 and Sar1^57^. Interestingly, for almost all COPII components (Sec16, Sar1, Sec24CD), disruption with two independent RNAi lines led to lethality of progeny, making memory analysis impossible. Only one Sec16 RNAi line survived, but this line showed no impact on learning (Figure S3F). This suggests that the COPII pathway (and transport of GPCRs generally) is required in MBN (or elsewhere with R13F02-gal4 expression) for survival, for unclear reasons. Sec24AB is not required in the MBN for survival, and perhaps has a specific role in transporting learning related GPCRs like R1.

If these proteins traffic R1 to regulate learning, then we anticipate that disruption of these trafficking proteins would alter other R1-mediated behaviors. Recently, it was shown that hungry flies have increased attraction to novel odors, and this requires R1 in MBN synapses^31^. R1 signals via distinct adenylyl cyclases resulting in distinct types of synaptic plasticity compared to aversive learning. In this assay, adult animals are either given water only (starved) or food (fed) for 50 hrs and then tested for their preference between a novel odor (MCH) and no odor (Figure 3G).

As shown previously, normally fed flies (with or without R1 disruption) avoid novel odors at the concentrations used in this assay (Figure 3H). This avoidance causes flies to have large negative odor attraction values. However, when control flies are starved, they become significantly less averse (more attracted) to the novel odor MCH, but if R1 is disrupted in the MBN, this significant shift in odor attraction does not occur, as reported previously^31^. Remarkably and similar to R1, disruption of Krz and CG13887 in the MBN prevented starvation-induced shift in odor attraction, but not Sec24AB (Figure 3I). Altogether, we have identified three trafficking genes orthologous to members of our Dop1R1 proximal proteome that regulate two distinct R1-mediated pathways and behaviors (Krz and CG13887), or just learning (Sec24AB).

### Trafficking proteins Sec24AB, CG13887, and Krz regulate DA signaling in MBN

To test whether these trafficking proteins that modulate R1-based behaviors alter the overall levels of R1 in the MBN, we imaged and immunostained brains expressing Dop1R1 endogenously tagged with mVenus^58^ and co-expressing RNAi for Sec24AB, Krz, CG13887, or no RNAi control, in the MBNs, and quantified the levels of R1. We found that the R1 abundance in the mushroom body lobes, as well as the fan-shaped body and central brain overall, was not affected by the MBN expression of RNAi for Sec24AB, Krz, or CG13887 (Figure 4A, B). These data indicate that the general production and transport of R1 out to the synaptic neuropil is not affected by these trafficking proteins. However, short-range trafficking of R1 to and from the plasma membrane at the synapse would not be measurable at this gross anatomical level.

**Figure 4.**
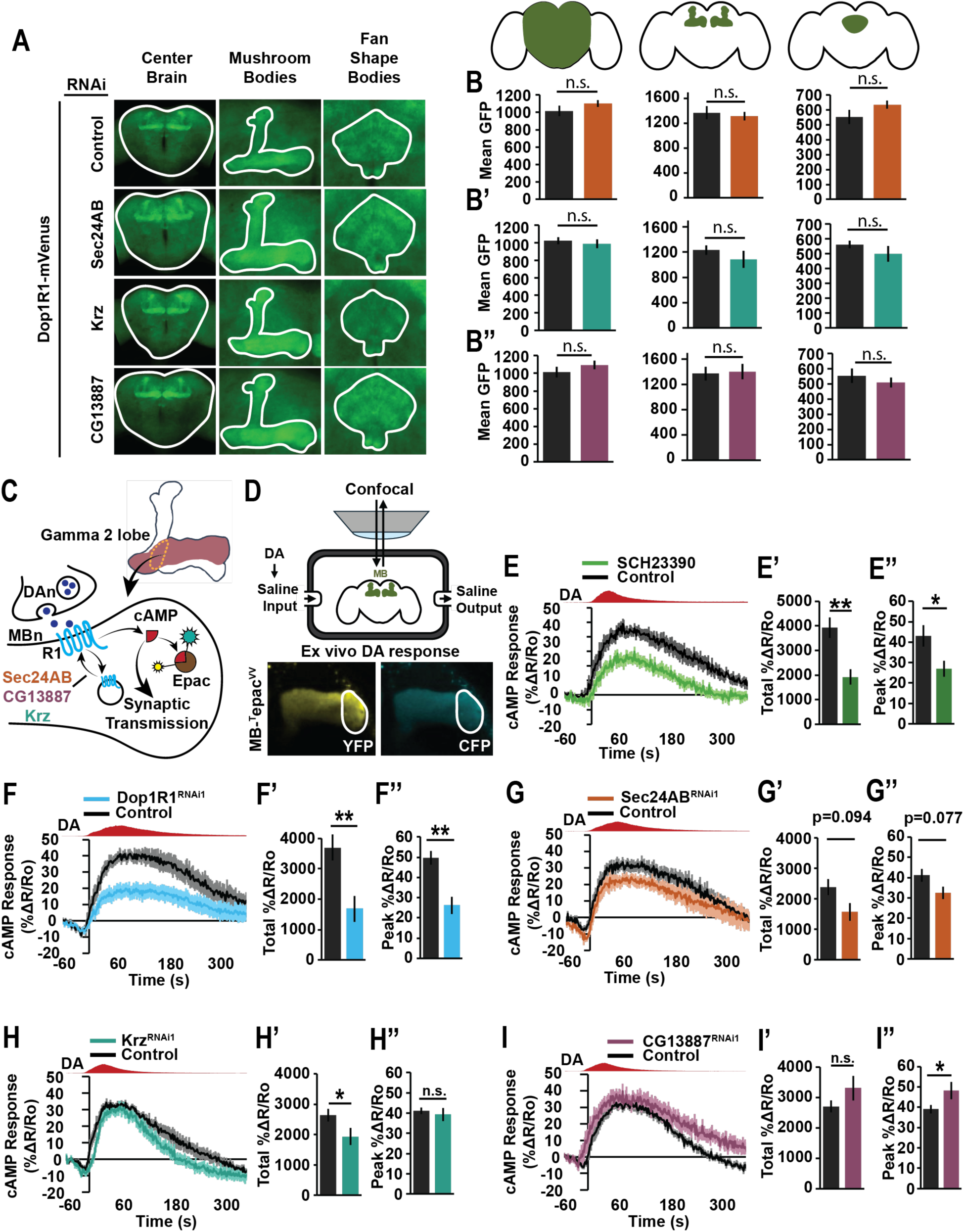
Trafficking proteins are essential for dopamine-dependent cAMP response to dopamine. (A) Representative images of the central brain, mushroom body, and fan shape body of control flies compared to flies with sec24AB, krz, and CG13887 RNAi in the MBn with endogenous Dop1R1-mVenus. (B-B’’) sec24AB, krz, and CG13887 RNAi in the MBn did not change the level of Dop1R1 in the center brain or the MB lobes compared to control brains (N=9-13, Students t-test). (C) Schematic representation of R1 signaling through cAMP in the gamma 2 MBn using MB-^T^epac^vv^ as a reporter. (D) Top: Schematic illustrating the experimental setup of the *ex vivo* imaging paradigm. Bottom: Representative images of the gamma 2 MBn lobe (White outline) showing the cAMP reporter MB-^T^epac^vv^ during the *ex vivo* imaging paradigm. (E) Treatment of brains with a D1-like receptor inhibitor significantly decreased cAMP response to 10mM DA perfusion. (E’) A sum of the total cAMP response from dopamine onset to the end of the recording, and (E’’) shows the average peak cAMP during the dopamine window. cAMP response is represented by a ΔR/Ro where R = ratio of CFP/YFP fluorescence intensity of cAMP reporter Tepac^vv^ and Ro = the average R during 10 frames of baseline recording (N=9. Mann-Whitney U test with Bonferroni correction for multiple comparisons **=P<0.01 and ***=P<0.001). (F) Dop1R1 knockdown causes a significant decrease in the cAMP response to 10 µM dopamine perfusion across the brain *ex vivo*. (N=9. Mann-Whitney U test with Bonferroni correction for multiple comparisons **=P<0.01 and ***=P<0.001). (G-G’’) Sec24AB knockdown decreases cAMP response in the gamma 2 lobe in response to 10mM dopamine perfused across the brain *ex vivo*. (N=9. Mann-Whitney U test with Bonferroni correction for multiple comparisons **=P<0.01 and ***=P<0.001). (H-H’’) CG13887 knockdown does not impact the mean cAMP response in the gamma 2 lobe in response to 10 mM dopamine perfused across the brain *ex vivo*. The peak response during dopamine is significantly increased (N=9. Mann-Whitney U test with Bonferroni correction for multiple comparisons #=P<0.1). (I-I’’) krz knockdown significantly decreases the mean cAMP response in the gamma 2 lobe, but did not impact the peak cAMP levels in response to 10mM dopamine perfused across the brain *ex vivo* (N=9. Mann-Whitney U test with Bonferroni correction for multiple comparisons #=P<0.1).

To more directly test the role of these trafficking proteins on DA->R1 mediated synaptic signaling in the MBN, we conducted *ex vivo* imaging of downstream cAMP response to DA perfusion across the brain using MBN driven cAMP FRET sensor (MB-Epac)^59^. We reasoned that if the disruption of these trafficking proteins altered the presence of R1 at the membrane, we should see altered DA-mediated cAMP in the MBN neuropil (Figure 4C). We measured cAMP responses (change in FRET ratio YFP/CFP compared to baseline) to a 30 sec perfusion of 10mM DA at the γ2 compartment of the MBN, a synaptic neuropil critical in the acquisition and expression of short-term aversive memory^27,60–64^ and starved odor avoidance behavior^31^ (Figure 4D). As expected, direct disruption of R1 signaling through treatment with the D1-like receptor inhibitor SCH23390, or R1 RNAi expression in the MBN, significantly impaired DA-mediated cAMP responses (Figure 4E, F) in terms of total (Figure 4E’, F’) and peak responses (Figure 4E’’, F’’). These experiments confirm that cAMP response to DA is highly dependent on R1.

Finally, we disrupted Sec24AB, Krz, and CG13887 trafficking proteins in the MBN and found that Sec24AB RNAi expression caused a trending, but not significant, disruption in overall and peak cAMP responses to DA (Figure 4G-G”), whereas, Krz RNAi expression significantly reduced total cAMP responses to DA, but not the peak (Figure 4H-H’’). On the other hand, CG13887 RNAi expression in the MBN significantly enhanced peak cAMP responses to DA, but not the total (Figure 4I-I”). These physiological data suggest that Sec24AB and Krz are required for high levels of cAMP in response to DA, whereas CG13887 normally suppresses cAMP response to DA, thus mirroring their respective effects on learning (Figure 3D-F).

## Discussion

For the first time, our study utilizes live-cell proximity labelling to canvas the distinct environments of two D1-like DA receptors, Dop1R1 and Dop1R2 (Figure 1), both critical regulators of distinct types of synaptic plasticity and memory processes in the MBs^2,8,27,28^. Drawing on reproducible results across two independent data sets, our data supports important and interesting insights regarding dopamine receptor localization, desensitization pathway interaction, and re-location after ligand binding, and potential distinctions between these two important receptors (Figure 2). Using our Dop1R proximal network, we employed an RNAi based gene disruption screen in fruit flies to identify DA receptor trafficking proteins involved in memory, in particular learning. We identified 3 genes (Sec24AB, Krz, and CG13887) that regulate two R1-mediated behaviors, memory acquisition and starvation induced odor attraction (Figure 3), and alter cAMP signaling in response to DA in MBN (Figure 4), likely through altered R1 receptor trafficking and presence at the surface of synapses. Furthermore, our data sets provide a foundation for future targeted disruption screens to study regulators of DA-mediated behaviors beyond just learning.

To begin, we comprehensively evaluated if protein fusion to either terminus of R1 and R2 dopamine receptors, as well as the presence of a linker, would alter live cell DA-receptor signaling, and found that N-terminal fusions of a SNAP protein dramatically reduced cellular responses to DA (Figure S1). We found that C-terminal fusion of SNAP along with GGS_4_ linker best preserved both receptors signaling. Replacing SNAP with the larger Turbo-V5 still preserved R1-cAMP signaling, but abrogated R2-Ca^2+^ signaling. We found that R2-Turbo-V5 expression in cells is reduced compared to PM and R1-Turbo-V5, given the same vector mass during transfection (Figure S1C); thus, we reasoned that the reduced R2 signaling (Figure 1C) is likely due to decreased receptor levels rather than altered functionality at the membrane. We adjusted DNA mass during transfection to achieve similar biotin labelling (Figure S1D) to normalize ligase and proteome levels to address this. In further support of R1 and R2 function being intact, we found Arrb2 was enriched to a similar level for both R1 and R2 only when DA was also present for both data sets (Figure 2K), suggesting R2 retains function at the plasma membrane despite the Turbo-V5 fusion. Thus, using independent TMT and label-free data sets (Figure S2), we have produced the first rich characterization of R1 and R2 cellular environments and potential interactors. While these proteomes are human proteins, there is remarkable conservation of function across the animal kingdom in terms of DA receptors^9,25^, downstream GPCR signaling elements, and cellular systems^65–67^. Given DA plays a vital role in a variety of behaviors and synaptic modulation via these two receptors^8,27,68–72^, we conclude that our data set will serve as a strong and relevant resource for behavior and synaptic biology researchers in the future.

Many GPCRs, including DA receptors in mammals, are located in lipid rafts in the plasma membrane where various signaling components are concentrated along with GPI-anchored proteins, heterotrimeric G proteins, and adenylyl cyclases to create signaling microdomains^73,74^. Our data indicates that in the basal condition, R1 is localized at the plasma membrane in lipid rafts along with GPI-anchored and lipid targeted proteins SMPDL3B, PRNP, CD44, and CD55, as well as GPCR signaling proteins like adenylyl cyclase ADCY5, and gamma subunit GNG12 as these proteins decrease in abundance in R1 samples, but not R2, once DA is applied (Figure 2L, left & S2I). On the other hand, our data suggests R2 is not located in lipid rafts, but close to proteins annotated for cell-cell adhesion and postsynaptic density as these terms were significantly enriched with an R2 bias (Figure 2H) and also show a basal bias for R2 (Figure 2L, right). We note that the top 2 post-synaptic density proteins with R2 bias were Ephrin type-B receptor 2 (Ephb2) and neuroplastin (Nptn) (Figure S2H). Both Ephrin signaling and Nptn mediate cell-cell signaling that play a role in synaptic plasticity underlying memory in mammals and *Drosophila*^45–48^. Interestingly, ephrin signaling was recently shown to regulate forgetting of labile memory in MBN similar to R2 itself^48^, indicating R2 and EPHB2 might interact or collaborate in active forgetting. Future studies are needed to firmly tie R2 function to cell-cell adhesion signaling in the MBN.

Our study indicates R1 and R2 have distinct differences once activated by DA. R1 internalization likely proceeds in a canonical manner in which it recruits the GPCR desensitization pathway including interaction and phosphorylation by GRK2,3, and subsequent recruitment of ARRB2 (Figure 2M & S2J)^5,51^. On the other hand, R2 only recruits ARRB2 after DA application for both data sets (Figure 2M & S2J), but is proximal to GRK2,3 in both basal and DA condition (Figure S2J). These results indicate R2 is desensitized in a non-canonical manner where ARRB2 is recruited independent of phosphorylation. In fact, in HEK293 cells, ARRB2 can still be recruited to D2-like DA receptors when all GRKs are removed^75^. The identity of this GRK independent recruitment is unknown and the consequences of GRK2/3 recruitment around R2 in a basal state remains unclear.

After recruiting ARRB2 and GRKs, activated receptors are usually internalized via clathrin-mediated endocytosis^5,44^. Both our data sets provide evidence that R1 is internalized via a clathrin based mechanism including close proximity to clathrin based endocytosis regulating proteins FNBP1, SYNRG, EPS15, and EPS15L1 (Figure 2C,G,I & S2G) and these pathways are biased to R1 over R2. Our data indicates ARRB2 was recruited to R2 when DA was present (Figure 2K,M, S2J) and early endosome membrane GO term was significantly enriched for both R1 and R2 with a DA bias (Figure 2L). Thus, while both R1 and R2 recruit ARRB2 after activation and are internalized to the early endosome, R1 is endocytosed via a clathrin based mechanism whereas it remains an open question what route R2 takes from the plasma membrane to the early endosomal system.

Both of our stringent statistical and no cutoff GO analysis revealed a close proximity of R1 to protein networks relating to the cytoskeleton (Figure 2E) which includes actin filament nucleation, depolymerization, and capping (Figure 2G,I). We also note that beyond actin, the spectrin cytoskeletal protein SPTAN1 is significantly more abundant in R1 samples compared to both PM and R2-Turbo samples (Figure 2B,C), and other spectrin members SPTBN1, SPTBN2, and SPTB are similarly R1-biased but not significant (Figures 2I, S2G). Furthermore, WASH complex components WASHC2A and WASHC2C are involved in actin mediated transport of vesicles in the endosome and recycling back to the plasma membrane^76^, and are R1-biased (Figure 2C,I & S2G). Thus, R1’s close association with cytoskeleton (Figure 2F) is likely due to the cytoskeleton’s critical role in clathrin-mediated endocytosis^44,77^, and endosomal sorting and recycling to the plasma membrane^78^. On the other hand, our evidence suggests R2 might not be recycled or trafficked in the endosome in the same manner.

We were able to capture a statistically significant Dop1R1/2 proximal proteome that spans much of the cellular compartments along the lifespan of a GPCR (Figure 2D,E) and most of these were common to both receptors including systems for transport from the ER to Golgi. We utilized this smaller set of proteins to interrogate the role of orthologous proteins in the *Drosophila* MBN for learning, and found two *Drosophila* proteins (Sec24AB, CG13887) orthologous to protein involved in ER-Golgi transport (Sec24B, BCAP29/31) regulate learning (Figure 3C-F & S3D). Sec24AB is a key component of the COPII transport vesicle, important for recognizing cargo^79^. While some parts of the COPII trafficking pathway are essential for its function as a whole, such as Sar1 and Sec16^80,81^, other proteins could be playing more specific roles. Expressing RNAi for essential COPII proteins in the MBN resulted in lethality in the animals, indicating their importance in general cell function. Sec24AB and Sec24CD play a role in recognizing specific signal sequences on transmembrane proteins, such as GPCRs, and preparing them for exit from the ER^79^. While Sec24CD RNAi expression in the MBN caused lethality, Sec24AB RNAi expression did not and only affected memory acquisition. This could indicate that Sec24AB transports cargo, like dopamine receptors, that play more specialized roles, whereas Sec24CD transports essential cargo. While disruption of Sec24AB in the MBN did not alter overall receptor level expression in the axonal projections (Figure 4B), it significantly reduced DA mediated cAMP in a MBN compartment that is critical for short-term memory (Figure 4G). We propose that reducing Sec24AB in turn reduces transport of R1 to the plasma membrane, and this reduced level of R1 at the surface abrogates the overall cAMP signaling during learning, leading to impaired memory acquisition. Future experiments are needed to identify and disrupt putative Sec24AB cargo sequences in the tail of R1 to validate this trafficking.

CG13887 is an uncharacterized protein that shares some homology with BCAP31. BCAP31 is an important trafficking protein in the ER serving several functions including recognizing and targeting abnormally folded proteins for degradation (i.e. quality control), and as a cargo receptor that helps export transmembrane proteins^82^. Loss of function of BCAP31 in humans is associated with an X-linked intellectual disorder, Mendelian syndrome^83–85^. We report here that CG13887 functions as a memory suppressor gene^56^ as its disruption in MBN increases the capacity for memory acquisition (Figure 3C,F,F’ & S3D bottom), likely through increased cAMP responses to DA (Figure 4) during learning. If CG13887 functions as a cargo receptor for R1 export from the ER, similar to Sec24AB, it is unclear why loss of this cargo receptor would increase R1->cAMP signaling. An alternative explanation is that CG13887, similar to BCAP31’s other role as a chaperone protein and quality control, functions to slow down R1 exit from the ER to aid proper folding. In this case, reduction of CG13887 chaperone would increase the rate of R1 exit from the ER and could increase the number of R1 molecules at the surface, albeit increasing the likelihood of aberrant R1 conformation. Finally, it is possible that CG13887 plays a novel role outside of those characterized by BCAP31 and future studies are needed to elucidate the mechanism of enhanced memory acquisition.

Both Turbo proteomics data sets demonstrate that human ARRB2 becomes proximal to either R1 and R2 only when DA is present (Figure 2K,M & S2J), indicating R1 and R2 activation recruits this arrestin. Here we targeted the only two *Drosophila* non-visual arrestins in the MBN, Krz (high rank for sequence homology) and CG32683 (moderate rank for sequence homology). We found that disruption of Krz, but not CG32683, significantly reduced maximal memory acquisition (Figure 3C,E & S3D,E). Furthermore, disruption of Krz in the MBN reduced cAMP responses to DA (Figure 4H), thus we have identified Krz as the arrestin that likely regulates R1 internalization in the MBN memory center. Given the role of Arrb2 in receptor activation-based desensitization, we might expect Krz disruption would lead to more R1 at the membrane due to reduced internalization. However, we see Krz disruption causes decreased cAMP responses to DA. We reason that constitutive Krz disruption could lead to long-term high levels of R1 that might lead to compensatory mechanisms to transport less R1 to the membrane leading to the reduced DA mediated cAMP we see.

Finally, R1 in the MBN plays a critical role in two adaptive behaviors, associative learning^2^ and starvation induced odor attraction (Figure 3H)^31^. Here we tested the role of trafficking proteins Sec24AB, Krz, and CG13887 in both behaviors and found that Krz and CG13887 play a role in both R1 mediated behaviors, whereas Sec24AB was specific to just learning (Figure 3C-F & 3I). This is further evidence that Krz and CG13887 indeed regulate R1 signaling. Interestingly, disruption of Krz and CG13887 have opposite effects on learning, but the same effect on starvation induced odor attraction. While the mechanistic reason for Sec24AB’s role only in learning, and Krz/CG13887 differential phenotypes across these two behaviors remains unclear, the temporal scale of DA->R1 signaling during learning is in seconds, whereas starvation induced attraction to novel odors likely involved DA->R1 signaling over 50 hours. This difference in timescales of signaling likely creates distinct demands on R1 receptor trafficking that impact these two behaviors differentially.

## Author contributions

J.A.B. planned all of the proteomics experiments, performed all of the proteomics experiments and analysis, and helped plan and oversee all behavioral and imaging experiments. R.L.D. helped with planning proteomics experiments and contributed to the interpretation of data. D.C.G planned all behavioral and brain imaging experiments, performed primary and secondary screening on candidates, followed up on candidates, and performed and analyzed imaging data. B.K. helped collect primary screen data. R.D. and R.E. performed follow-up experiments on candidates that passed screens. J.A.B., D.C.G., and R.L.D. wrote the manuscript.

## Acknowledgement

Research in the laboratory of R.L.D. is supported by NIH grant R35NS097224. Research in the laboratory of J.A.B. is supported by Natural Sciences and Engineering Research Council grant RGPIN-2021-02545. This work is supported in part by The Herbert Wertheim UF Scripps Institute for Biomedical Innovation & Technology Mass Spectrometry and Proteomics Core, with particular thanks to George Tsaprailis and Gogce Crynen for performing mass spectrometry and analysis on protein samples and for providing helpful discussion and advice.

## Data availability statement

The mass spectrometry proteomics data sets have been deposited to MassIVE:

1. Label free data set - MSV000101396 (password: xkEk20Bci53J9k5S) https://massive.ucsd.edu/ProteoSAFe/dataset.jsp?task=270c5107fa634230936eadf6ab714d8e
2. TMT data set - MSV000101435 (password: YPOl9Y709yfrsTcP) https://massive.ucsd.edu/ProteoSAFe/dataset.jsp?task=016c873559cd48daa6f4c7c5560a66d4

## Supporting information

The following supporting information is available free of charge at ACS website http://pubs.acs.org

1. Figure S1. Verification of receptor function based on tag placement and confirmation of biotinylation in cell culture
2. Figure S2. Comparison of TMT and label-free data sets by protein abundance and GO analysis
3. Figure S3. Further characterization of behavioral phenotypes for Sec24AB, Krz, and CG31887.
4. Table S1. Fly lines used for behavioral and physiological experiments.
5. Supporting information for experimental procedures
6. Supporting spreadsheet 1 - Protein level quantitation, comparisons, and statistics
7. Supporting spreadsheet 2 - Gene ontology and Pathway enrichment and statistics

