## Supporting information for "Proximity labelling of D1-like dopamine receptors reveals distinct cellular environments and uncovers trafficking proteins that regulate DA mediated behaviors in *Drosophila*"

130 Scripps Way

Jupiter, Florida, 33458

Table of Contents:

1. Figure S1. Evaluation of tag placement, biotinylation, and expression of Turbo constructs in cells with or without biotin and with or without Turbo-V5.

2. Figure S2. Comparison of TMT and Label-free data sets, and protein networks from enriched GO terms.

3. Figure S3. Trafficking proteins play a role in regulating memory acquisition.

4. Table S1. Fly lines used for behavioural and physiological experiments.

5. Supplemental information for experimental procedures.

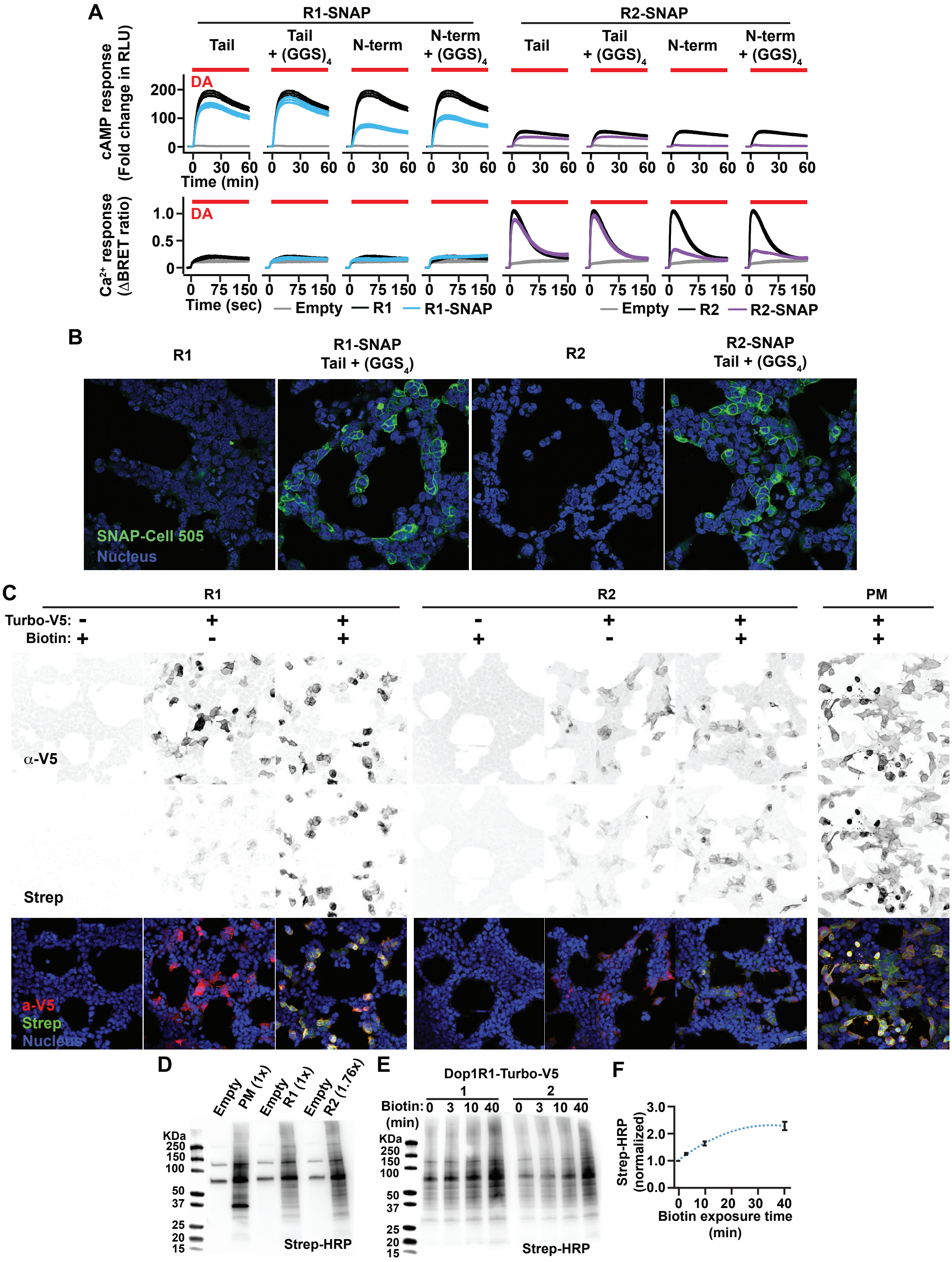
A
R1-SNAP
R2-SNAP
Tail
Tail + (GGS)4
N-term
N-term + (GGS)4
DA
cAMP response (Fold change in RLU)
Time (min)
Ca2+ response (ΔBRET ratio)
Time (sec)
Empty
R1
R1-SNAP
R2
R2-SNAP
B
R1
R1-SNAP Tail + (GGS4)
R2
R2-SNAP Tail + (GGS4)
SNAP-Cell 505
Nucleus
C
PM
Turbo-V5:
Biotin:
α-V5
Strep
a-V5
Strep
Nucleus
D
Empty
PM (1x)
R1 (1x)
R2 (1.76x)
KDa
Strep-HRP
E
Dop1R1-Turbo-V5
Biotin: (min)
Strep-HRP
F
Strep-HRP (normalized)
Biotin exposure time (min)

Figure S1

**Figure S1. Evaluation of tag placement, biotinylation, and expression of Turbo constructs in cells with or without biotin and with or without Turbo-V5.**

(A) Live cell imaging of cAMP (fold change in luminescence, top) and intracellular $Ca^{2+}$ (fold change in BRET ratio, bottom) responses to 100 μM DA in cells expressing empty vector, R1 (left), or R2 (right) with or without SNAP tag fused to different termini and with or without $GGS_4$ linker (n=6). (B) Expression of R1 or R2 with or without SNAP tag at the tail with a $GGS_4$ linker in cells visualized with a SNAP-Cell 505 ligand-fluorescent dye (green) at a cross-sectional plane. Cells counterstained for the Nucleus (blue). (C) Expression of R1, R2, or PM-Turbo-V5 in cells and Turbo labelling measured by anti-V5 and Strep-488 staining, respectively, with or without biotin (500μM, 1.5 hr) and Turbo-V5 fusion. a-V5 and STREP are maximum Z projections, and Nuclear stain is an average Z projection. Intensities were not adjusted to show reduced R2-Turbo expression compared to other constructs. (D) Representative western blot of Strep-HRP staining showing similar biotinylation levels between PM, R1, and R2-Turbo-V5 when transfected with adjusted DNA mass (PM:R1:R2, 1:1:1.76). (E) Representative western blot of Strep-HRP staining showing biotinylated proteomes from cells exposed to 500μM biotin media for 0, 3, 10, and 40 min. (F) Quantitation from 3 biological replicate western blots of Strep-HRP staining as a function of biotin exposure time, normalized to no biotin (0 min). The blue dotted line is a polynomial fit to the data.

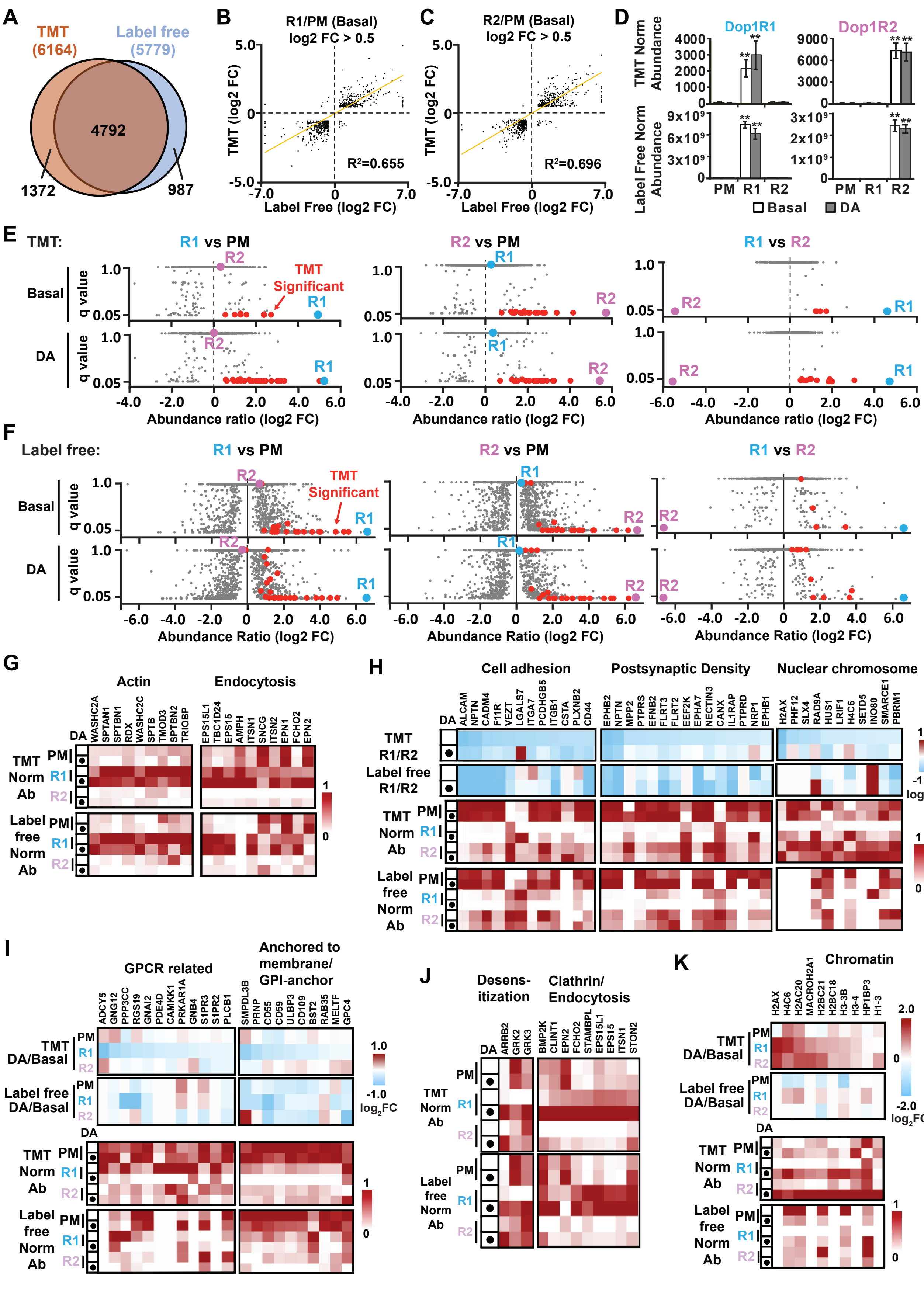
A
TMT (6164)
Label free (5779)
4792
1372
987
B
R1/PM (Basal) log2 FC > 0.5
TMT (log2 FC)
Label Free (log2 FC)
R²=0.655
C
R2/PM (Basal) log2 FC > 0.5
R²=0.696
D
Dop1R1
Dop1R2
TMT Norm Abundance
Label Free Norm Abundance
PM R1 R2
Basal DA
E
TMT:
R1 vs PM
R2 vs PM
R1 vs R2
TMT Significant
Basal
DA
q value
Abundance ratio (log2 FC)
F
Label free:
Abundance Ratio (log2 FC)
G
Actin
Endocytosis
H
Cell adhesion
Postsynaptic Density
Nuclear chromosome
I
GPCR related
Anchored to membrane/ GPI-anchor
J
Desens-itization
Clathrin/ Endocytosis
K
Chromatin

Figure S2

**Figure S2. Comparison of TMT and Label-free data sets, and protein networks from enriched GO terms, related to Figure 2.**

(A) Venn diagram of proteins quantitated in TMT and Label-free data sets. (B) Average abundance ratio (R1 Basal/PM Basal) for TMT proteome as a function of this ratio for the Label-free proteome on proteins with log2 ratio > 0.5. Yellow line indicates a linear regression fit and corresponding $R^2$ value is shown. (C) Same as (B), but using R2 Basal/PM Basal ratios. (D) Normalized abundance for Dop1R1 (left) and Dop1R2 (right) measured across basal and DA conditions for PM, R1, and R2-Turbo-V5 samples for TMT and label-free data sets. (E) Volcano plots showing the q-value as a function of abundance ratios for R1/PM (left), R2/PM (middle), and R1/R2 (right) for Basal (top) and DA (bottom) conditions for the TMT data set. R1, R2, and significant proteins (q<0.05, Figure 2B,C) are highlighted in color. (F) Same as (E) but for the Label-free data set. R1 and R2 are highlighted in color. Orange color are proteins that were significant (q<0.05, Figure 2B,C) in the TMT data set. (G) Heatmap of normalized abundance for proteins shown in Figure 2I, belonging to the top representative GO terms for actin and endocytosis for both TMT and label-free data sets. Abundance is normalized across all conditions/samples on a scale of 0 to 1 for each protein. (H) Heatmap of abundance ratios (R1/R2) (top) and normalized abundance (bottom) for proteins belonging to top representative GO terms (Figure 2H) with an R2 bias for both TMT and label-free data sets. (I) Heatmap of abundance ratios (DA/Basal) (top) and normalized abundance (bottom) for proteins with the strongest Basal bias (lowest DA/Basal ratio) belonging to the top representative and significant GO terms (Figure 2L, left) for both TMT and label-free data sets. (J) Heatmap of normalized abundance for proteins with the strongest DA bias (highest DA/Basal ratio) belonging to the top representative and significant GO terms (Figure 2L,2M) for both TMT and label-free data sets. (K) Heatmap of abundance ratios (DA/Basal) (top) and normalized abundance (bottom) for proteins with DA bias belonging to the significantly enriched Chromatin GO term (Figure 2L, right) for both TMT and label-free data sets.

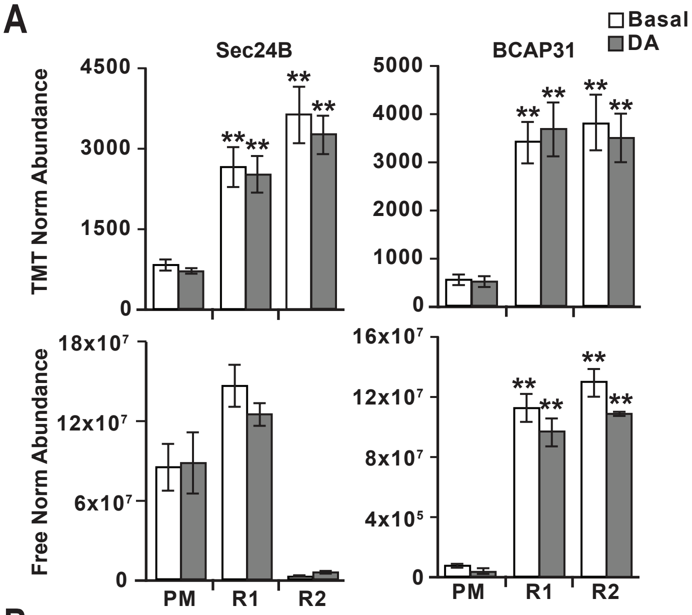


B

| | OCT avoidance | P-value | MCH avoidance | p-value |
|---|---|---|---|---|
| Control (krz) | 0.592 +/- 0.09 | | 0.798 +/- 0.05 | |
| Krz | 0.580 +/- 0.06 | 0.5803 | 0.598 +/- 0.05 | 0.0479 |
| Control (Sec24AB) | 0.597 +/- 0.04 | | 0.569 +/- 0.03 | |
| Sec24AB | 0.599 +/- 0.06 | 0.97062 | 0.600 +/- 0.06 | 0.531503 |
| Control (CG13887) | 0.712 +/- 0.05 | | 0.524 +/- 0.06 | |
| CG13887 | 0.710 +/- 0.07 | 0.55097 | 0.420 +/- 0.1 | 0.439547 |

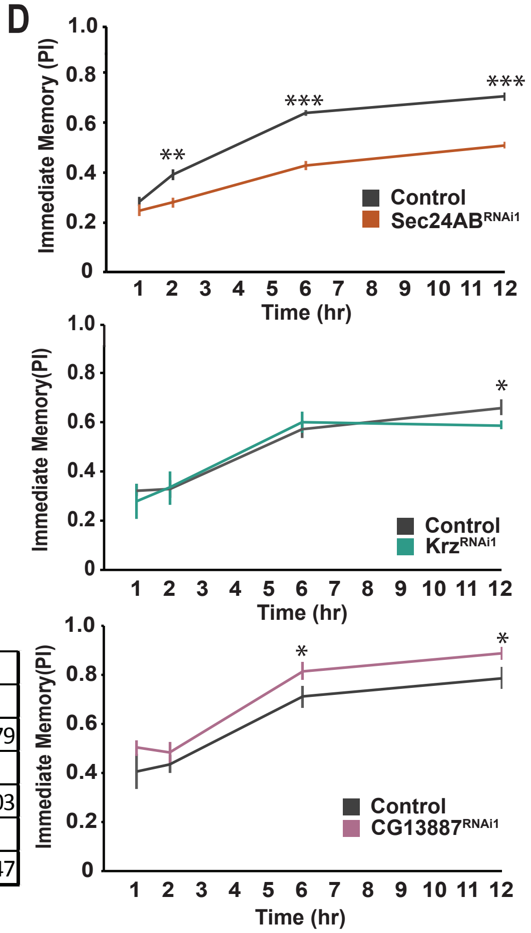


C

| | Shock Avoidance | P-value |
|---|---|---|
| Control (krz) | 0.788 +/- 0.03 | |
| Krz | 0.819 +/- 0.05 | 0.615 |
| Control (Sec24AB) | 0.711 +/- 0.05 | |
| Sec24AB | 0.711 +/- 0.02 | 0.997 |
| Control (CG13887) | 0.706 +/- 0.03 | |
| CG13887 | 0.710 +/- 0.04 | 0.925 |

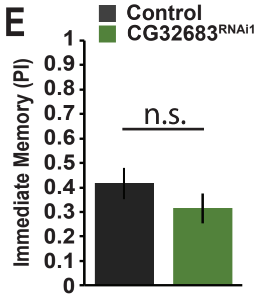


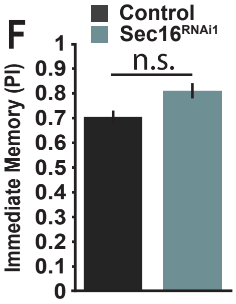


**Figure S3**

**Figure S3. Trafficking proteins play a role in regulating memory acquisition.**

(A) Normalized abundance for Human Sec24B (left) and BCAP31 (right) measured across basal and DA conditions for PM, R1, and R2-Turbo-V5 samples for TMT and label-free data sets. (B) Expression of RNAi against *Sec24AB*, *krz*, and *CG13887* did not alter innate odour response to OCT, but *krz* RNAi did have a reduced avoidance response to MCH compared to controls, while *Sec24AB* and *CG13887* RNAi did not (N=8, Student t-test). (C) Expression of RNAi against *Sec24AB, krz*, and *CG13887* did not alter the flies shock response to 90V electric shock (N=8, Student t-test). (D) Expression of *Sec24AB* RNAi in the MBn decreases memory acquisition when flies were trained with 2, 6, and 12 shocks (Top). Expression of *krz* RNAi in the MBn decreases memory acquisition when flies were trained with 12 shocks (Middle). Expression of *CG13887* RNAi in the MBn increases memory acquisition when flies were trained with 6 and 12 shocks (Bottom). Memory acquisition was tested by varying the number of shocks given during training and testing immediate memory following training. Control: R13F02>+; UAS-Dicer (N=8. Students t-test, * = P<0.05, **= P<0.01, ***=P<0.001). (E) Expression of CG32683 RNAi in the MBn did not alter learning in flies compared to control (N=8, Students t-test). (F) Expression of Sec16 RNAi in the MBn did not alter learning in flies compared to control (N=8, Students t-test).

**Supporting information for experimental procedures**

**Cell culture**

Cells were grown in “normal culture media” (see Supporting information) that includes Dulbecco’s modified Eagle’s medium (DPBS) supplemented with 10% fetal bovine serum, MEM non-essential amino acids (Life Technologies), 1 mM sodium pyruvate, and 100 units/ml penicillin and 100 µg/ml streptomycin at 37°C in a humidified incubator equilibrated with 5% $CO^2$. Cells were seeded onto 6 or 15 cm culture plates at 1.3x10^6/3 ml culture media, or 8.1x10^6/ 18.75 ml culture media for microscopy and proteomic experiments, respectively, and grown till 50-60% or 80-90% confluency for microscopy or cell signaling/proteomics, respectively.

**Live cell imaging procedures for cAMP and $Ca^{2+}$**

For cAMP, cells were detached with 1ml of CO2 independent medium (Invitrogen) containing 10% FBS, washed 2x, and resuspended in the same buffer ($2x10^6$ cells/ml). The GloSensor cAMP Reagent (Promega, #E1290) was prepared according to manufacturer’s instructions to make a cAMP reagent stock. Twenty-five ul of cells and 25 ul of 2X GloSensor cAMP Reagent (4% cAMP reagent stock in $CO_2$ independent media with 10% FBS) were added to each well of a 96-well plate and mixed, incubated for 2hr, and Luminescence (615 nm) was monitored with CLARIOstar microplate reader. Fifty µl of 100 µM DA dissolved in PBS containing 0.5 mM $MgCl_2$ and 0.1% glucose was applied to cells after a 30 sec baseline. cAMP responses were calculated as a fold change (FC) in luminescence (Relative Luminescence Units, RLU) from baseline

(RLU=L(t)/Baseline), where baseline was the average luminescence in the 30 sec prior to DA application. Sample size was 6 biological replicates (6 wells).

For $Ca^{2+}$/CalfluxVTN assays, transfected cells were washed once with PBS containing 0.5 mM MgCl2 and 0.1% glucose and detached by gentle, repeated pipetting in the same solution. Cells were harvested by centrifugation at 500xg for 5 min and resuspended in DPBS containing 0.5 mM MgCl2 and 0.1% glucose. Approximately 50,000 to 100,000 cells in 100 ul were added per well in 96-well flat-bottomed white microplates (Greiner Bio-One). The Nluc substrate, furimazine, was purchased from Promega and used according to the manufacturer's instructions. DA (100 μM) dissolved in PBS containing 0.5 mM MgCl2 and 0.1% glucose was applied (50 μl) after a 10 sec baseline. BRET measurements were made using a CLARIOstar microplate reader at room temperature. The BRET signal was calculated by measuring the ratio of light emitted by the Venus reporter (535 with 30 nm band pass filter) relative to light emitted by a Nluc reporter (475 nm with 30 nm band pass filter) every 0.4 seconds[38]. The fold change (FC) in BRET ratio across time was calculated by dividing the points by the average BRET ratio for 10 sec prior to DA injection (baseline). Sample size was 6 biological replicates (6 wells).

**V5, biotin, and SNAP tag staining and microscopy**

For SNAP tag staining (Figure S1B), 300,000 cells in 500 ul were seeded onto poly-L-lysine cover slips placed in 12 well plates and grown till ~50-60% confluency (to reduce density for better microscopy of single cells) and then transiently transfected (as above) with 1 ug/well of pCAG vectors containing Dop1R1 and Dop1R2 with or without $GGS_4$-

SNAP fused to the tail and empty vector control. After ~19 hrs post transfection, cells were treated with 30 min exposure to 3 μM of cell permeable SNAP ligand (SNAP-Cell 505-Star from NEB) that contains the BG binding group conjugated to green fluorophore. Excess SNAP ligand were then removed through the following wash/incubations with normal culture medium at 37$^o$C, 5% $CO_2$: 3x with 2 min incubations, 1x with 30 min incubations, and 3 times with 2 min incubations. To stain the nucleus, cells were incubated with normal culture media with 2 drops/ml of cell permeable NucBlue (Invitrogen, R37605), followed by two normal culture media washes for 2 min at 37$^o$C, 5% $CO_2$. Cells were then fixed by replacing media with 4% paraformaldehyde in DPBS for 20 min at room temperature, then washed 3 times with 2 min incubation in DPBS, prior to imaging via Leica SP8 confocal microscope at 1024x1024 using 488 nm and 740 nm multiphoton lasers and detectors set to 555-625 nm and 415-485 nm for SNAP-Cell-505 and NucBlue fluorescence, respectively.

For anti-V5 and biotin staining, cells were fixed with 4% paraformaldehyde in DPBS for 20 min at RT. After 3x DPBS washes, cells were permeabilized with ice cold methanol for 5 min at -20$^o$C. After 3x DPBS washes, cells were then incubated with primary antibody against V5 (1:100, chicken anti-V5 polyclonal, Abcam, ab9113) and 3% bovine serum albumin in DPBS for 1 hr at 4$^o$C. After 3x DPBS washes, cells were incubated with secondary antibody (1:500, goat anti-chicken IgG Alexa 633, Life Technologies, A21103), Streptavidin-Alexa 488 (1:200 from 2mg/ml stock, ThermoFisher, S11223), and 3% bovine serum albumin (BSA) in DPBS for 1 hr at 4$^o$C. After 3x DPBS washes, cells were incubated with NucBlue stain (2 drops/ml, Invitrogen, R37605) for 5 min. Finally, after 3x DPBS washes, coverslips were imaged via Leica

SP8 confocal microscope at 1024x1024 using 488 nm, 633 nm, and 740 nm multiphoton lasers and detectors set to 500-550 nm, 640-700 nm, and 418-485 for Streptavidin-Alexa 488, goat anti-chicken Alexa 633, and NucBlue fluorescence, respectively.

**Creating cell lysates for proteomics and western blot analysis**

After biotin and/or DA treatment, cells were first dissociated from the plates using pipette aspiration with DPBS that is ice cold to preserve cells and kill Turbo labelling. The remaining steps and reagents are kept on ice. To remove the excess biotin, we doubled cell suspension volume by adding DPBS and washed via pipetting up and down most of the volume 10x, prior to pelleting cells (1500 rpm, 4 min at 4$^{o}$C). Cells were homogenized/lysed by first resuspended in lysis buffer consisting of 50 mM Tris-HCl (pH 8.0), 150 mM NaCl, 0.1% SDS, 0.5% sodium deoxycholate, 1% Triton X-100, and EDTA-free protease inhibitor complex (Roche, 4693132001), and vigorously pipetting (without creating bubbles), and then fully homogenized via motorized pestle before obtaining the clarified supernatant (10,000 rpm at 4$^{o}$C). Protein concentrations across lysates in an experiment were normalized via BCA assay (Pierce BCA Protein Assay Kit, ThermoFisher, 23225), prior to western blot or bead incubation. For WB analysis, lysates were processed and western blots were made using standard protocols[33] with the following specifics. Twenty ug of protein lysates were ran in each lane of a 10 well SDS gel (4-20%) via electrophoresis prior to transferring proteins to PVDF membrane, and blocking with non-fat milk (5%) in Tris buffered saline with Tween-20 (0.1%) (TBST). Blots were then washed 5x (5 min) with TBST and incubated with Streptaviding-HRP (40 ng/ml, Sigma-Aldrich, OR03L) and 3% BSA in TBST for 1 hr at 4$^{o}$C, prior to 5x (5 min) washes with TBST and subsequent HRP substrate incubation and imaging.

**LC-MS/MS protein identification and quantification for Label free data set**

For the label free data set, eluted proteins (6 experimental conditions, 2-3 biological replicates/condition) were precipitated overnight with 4X ice cold acetone (v:v). Protein pellets were then solubilized in 100 µL 6M urea/50 mM Tris pH8, reduced using 3 µL of 0.5M DTT at 45$^{o}$C for 45 min and then alkylated in the dark at room temperature for 30 min using 6 µL of 0.5M IAA. Proteins were acetone precipitated overnight once more, solubilized in 50mM TEAB, and digested overnight with trypsin (1 µg) at 37 $^{o}$C.  A colorimetric peptide assay using the Pierce™ Quantitative Colorimetric Peptide Assay kit (Thermo Fisher Scientific, Waltham, MA) was performed according to the manufacturer's instructions to determine the peptide yield. LC-MS/MS analysis of peptides was carried out using an Orbitrap Fusion Tribrid mass spectrometer, following 2 µg capacity ZipTip (Millipore, Billerica, MA) C18 sample clean-up according to the manufacturer's instructions. Peptides were eluted from an Acclaim PepMap™ RSLC nano Viper analytical column (75-µm ID × 15 cm, Thermo Scientific, San Jose, CA) using a gradient of 5-25% solvent B (80/20 acetonitrile/water, 0.1% formic acid) in 180 min, followed by 25-44% solvent B in 60 min, 44-80% solvent B in 0.10 min, a 5 min hold of 80% solvent B, a return to 5% solvent B in 0.10 min, and finally a 20 min hold of solvent B. All flow rates were 300nL/min delivered using a nEasy-LC1000 nano liquid chromatography system (Thermo Fisher Scientific, San Jose, CA). Solvent A consisted of water and 0.1% formic acid. Ions were created at 1.9kV using the Nanospray Flex$^{TM}$ ion source (Thermo Scientific, San Jose, CA). Data dependent scanning was performed by the Xcalibur v 4.0.27.10 software using a survey scan at 120, 000 resolution in the Orbitrap analyzer scanning mass/charge (m/z) 380-2000 followed by higher-energy collisional dissociation (HCD) tandem mass spectrometry (MS/MS) at a normalized collision energy of 30% of the most intense ions at maximum speed, at an automatic gain control of 1.0E4. Precursor ions were selected by the monoisotopic precursor selection (MIPS) setting to peptide and MS/MS was performed on charged species of 2-8 charges at a resolution of 30,000. Dynamic exclusion was set to exclude ions after two times within a 30sec window, for 20sec. Quantitative analysis of the label free experiments was performed simultaneously to protein identification using Proteome Discoverer 2.4 software. The precursor and fragment ion mass tolerances were set to 10 ppm, 0.02Da, respectively), enzyme was Trypsin with a maximum of 2 missed cleavages and Uniprot Human proteome FASTA file (downloaded in 2017) with six additions (P00761|TRYP_PIG, P22629|SAV_STRAV, P02769|ALBU_BOVIN, P06709|BIRA_ECOLI, Q24563|DOPR2_DROM, P41596|DOPR1_DROME) was used in SEQUEST searches. The following settings were used to search the data; dynamic modifications; Oxidation / +15.995Da (M), Deamidated / +0.984 Da (N, Q), Biotin / +226.078 Da (K and N-terminus , Acetyl 42.0106 Da (protein N-terminus), and static modifications of Carbamidomethyl +57.021 (C). Only unique+ Razor peptides were considered for quantification purposes. *Percolator* feature of Proteome Discoverer 2.4 was used to set a false discovery rate (FDR) of 0.01. Co-isolation threshold and SPS Mass Matches threshold were set to 50 and 65, respectively. *Minora feature detector* node used with default settings. Total Peptide Amount normalization method was used to correct for loading bias. Average of Top 3 peptide precursor area was processed through *Protein Abundance Based* method to calculate the protein level ratios.

| Gene | Source | Identifier |
|---|---|---|
| 14-3-3zeta | VDRC | KK 104496 |
| 5-HT2A | BDSC | RRID:BDSC_56870 |
| 5-HT2B | BDSC | RRID:BDSC_60488 |
| Alpha-spec | BDSC | RRID:BDSC_56932 |
| Arll | BDSC | RRID:BDSC_27052 |
| CG11374 | BDSC | RRID:BDSC_61863 |
| CG13526 | VDRC | KK 108977 |
| CG13887 | VDRC | KK 106452 |
| CG13887 | VDRC | GD 23207 |
| CG32683 | VDRC | KK 104029 |
| CG32683 | VDRC | KK104029 |
| CG3309 | VDRC | KK 110363 |
| CG3408 | VDRC | GD 36306 |
| CG3708 | BDSC | RRID:BDSC_67336 |
| CG5335 | VDRC | KK 101295 |
| CG5823 | VDRC | KK 108292 |
| CG6766 | BDSC | RRID:BDSC_55745 |
| CG7945 | VDRC | KK 101355 |
| CG9920 | VDRC | KK 102856 |
| cindr | BDSC | RRID:BDSC_38976 |
| Cip4 | VDRC | KK 108625 |
| DopR1-Venus | Kondo et al., 2020 | |
| Eip63F-1 | VDRC | KK 102809 |
| Ero1L | VDRC | KK 110454 |
| FAM21 | VDRC | GD 23832 |
| FKbp14 | BDSC | RRID:BDSC_67224 |
| galectin | VDRC | KK 107054 |
| GD Control | VDRC | GD 60000 |
| Gdap2 | VDRC | KK 100622 |
| Gmap | BDSC | RRID:BDSC_64863 |
| His2Av | VDRC | KK 110598 |
| InaD | BDSC | RRID:BDSC_52313 |
| Jafrac1 | BDSC | RRID:BDSC_32498 |
| Jafrac2 | VDRC | KK 104006 |
| KK Control | VDRC | KK 60100 |
| Krz | VDRC | KK 103756 |
| Krz | VDRC | GD 41559 |
| MB-TEPACVV | Boto et al., 2014 | |
| mil | VDRC | KK 106545 |
| Moe | VDRC | KK 110654 |
| mRpL 19 | VDRC | GD 29474 |
| Pex19 | VDRC | KK 100746 |

| | | |
|---|---|---|
| Poly | BDSC | RRID:BDSC_65225 |
| Prx3 | BDSC | RRID:BDSC_60475 |
| R13F02-gal4 | BDSC | RRID:BDSC_48571 |
| Rbsu-5 | BDSC | RRID:BDSC_57459 |
| ReepB | BDSC | RRID:BDSC_62476 |
| Rhea | BDSC | RRID:BDSC_33913 |
| RhoGAP1A | VDRC | KK 105202 |
| Sar1 | VDRC | GD 34191 |
| Sar1 | VDRC | KK 108458 |
| Sar1 | BDSC | RRID:BDSC_32364 |
| Sec16 | VDRC | KK 109645 |
| Sec16 | BDSC | RRID:BDSC_53917 |
| Sec24AB | VDRC | KK 107154 |
| Sec24AB | VDRC | GD 44461 |
| SREBP | VDRC | GD 37641 |
| sty | VDRC | GD 6948 |
| tacc | VDRC | KK 101439 |
| Tace | VDRC | KK 106335 |
| Trip Control (2nd) | BDSC | RRID:BDSC_36304 |
| Trip Control (3rd) | BDSC | RRID:BDSC_36303 |
| Tx1 | VDRC | KK 110341 |
| UAS-Dicer2 | VDRC | 60008 |
| wts | VDRC | GD 9928 |

**Table S1. Fly lines used for learning screen and physiology experiments.**
